# Dual mechanical impact of β-escin on model lipid membranes

**DOI:** 10.1101/2023.06.15.545043

**Authors:** Lara H. Moleiro, María T. Martín-Romero, Diego Herráez-Aguilar, José A. Santiago, Niccolo Caselli, Carina Dargel, Ramsia Geisler, Thomas Hellweg, Francisco Monroy

## Abstract

Using artificially reconstituted membranes based on structurally liquidlike phospholipids, we have performed an experimental study on the mechanical impact of the saponin β-aescin, aka escin, a natural biosurfactant extracted from the seeds of the horse chestnut tree *Aesculus hippocastanum*. The paper focusses on the modulable interaction of escin with DMPC in model membranes in the form of bilayer vesicles and Langmuir monolayers. As regarding to their dual mechanical membrane behavior being both soft solids and viscoelastic fluids, we have outlined the principal energetic and kinetic features describing the insertion of escin as transversally adsorbed or longitudinally integrated within the model membranes. At connection with the structural phase behavior assessed by dedicated microscopies of surface fluorescence and Brewster angle reflectivity, these hybrid escin / phospholipid membranes have been revealed to possess dual mechanical properties connected to their structural rigidness and fluidity behaving both in one way and another. In particular, we observe a soft glassy rheology typical for liquid-crystalline ordered phases at low temperature, which turns into a fluidlike viscoelasticity characteristic of the disordered phases at high physiological temperature. These original results have been discussed in a physicochemical perspective that may pave new avenues of material engineering and / or pharmacological design exploiting the dual mechanical impact of escin as a mechanical modulator of the cellular membrane.

## 1. Introduction

The fluid mosaic paradigm structurally establishes the biological membranes with a mechanical resilience as flexible objects endowed of molecular mobility [1]. Such compositional mosaicity provides the embedded proteins with a functional diffusivity as laterally structured across membrane domains dispersed in a continuous liquid phase [2]. Whereas fluids yet being practically incompressible can flow under applied shear stress, solids however resist tightly to shear deformations, so that, biological membranes require structural rigidity and lateral fluidity as being solid and liquid both i.e., two ambivalent mechanical aspects comprised all in one requested for a functional homeostasis [3–5]. Two prior pivotal studies have hypothesized such mechanical duality as necessarily stemming from a functional membrane viscoelasticity endowed by the heterogeneous composition of the monolayer leaflets [6,7]. Such dual viscoelasticity typical of biological membranes recapitulates the high dynamic viscosity required enough high for regulated fluidity together with a finite shear modulus that imparts sufficient structural resistance [8], being both engendered by the liquid crystalline arrangement of the membrane lipids [9–12]. The dual viscoelastic hypothesis of biological membranes above posed with artificial models has been further substantiated in a wealth of experimental studies with real living systems [13–17]. Those previous works have evidenced a variety of mechanical membrane behaviors emerged under biochemical regulation of the lipid flows, for instance, metabolic in-membrane conversions of fluid sphingomyelins into solid ceramide leading to lateral tension gradients [13,14] -as far as they elicit transformations of cell shape [18], and a functional capacity to support propagating mechanical signals -as a relevant transduction apparatus in the cellular mechanobiology [15,19]. As a matter of fact, the membrane viscoelastic stresses have been also revealed with a crucial role in the development of cancer [16,20], and in the involvement of membrane mechanics for cell hereditariness [17,21]. Despite the advances in understanding the functional dynamics of biomembranes with experimental models of ordering lipid mosaicity [7-13,23-29], and the very detailed theoretical approaches relating to their liquid-crystalline viscoelasticity [30-38], the mechanical paradigm of biological membranes is still incomplete as far as the impacts of ordering-imparters on the membrane stresses remain unidentified hitherto. Molecular membrane insertion may eventually occur either as an intrinsic compositional integration from the metabolic lipid factories (longitudinal lipogenesis), or as an extrinsic incorporation from the membrane environment (transverse lipid traffic). In this work, we report a comparative rheological study on the dual mechanical impact of intrinsic (longitudinal) vs. extrinsic (transverse) lipid insertion in membrane models constituted by the phospholipid dimyristoylphosphatidylcholine (DMPC). Their surface rheology will be studied under dually vectorized (longitudinal / transverse) insertion of β-aescin, a saponin biosurfactant mimicking the effect of solidlike stresses as imparting enhanced membrane stiffness and additional viscous friction [39,40].

Saponins are a family of natural triterpenoids obtained as plant extracts that endow emulsifier and foaming activities in aqueous media. Their biochemical classification as biosurfactants make them useful for technological development of functional foods, cosmetics and drugs. Some saponins also exhibit cytotoxic effects on cancer cells [41-43], however, their systemic propensity to induce hemolysis limits their anti-tumorigenic potential in therapy. Particularly, the amphiphilic molecule β-aescin, aka escin, is the major component isolated as a natural product on the mixture of saponins extracted from the seed of the chestnut obtained of *Aesculus hippocastanum*. This derivative of the glucopyranosic acid has a chemical formula as drawn in Figure 1A. The amphiphilic molecular structure is clearly discernible as an oligosaccharide subsystem in the polar head linked to a triterpene core ring plane subsystem –the aglycone constituting the hydrophobic moiety (see Fig. 1A; inset). Some anti-inflammatory, anti-edematous and venotonic effects are attributed to this biosurfactant [44], especially in treatments of chronic venous insufficiency (CVI) [45-47]. The etiology of CVI is very complex and involves genetic susceptibility and environmental factors leading to several inflammatory pathways that become activated to cause vein wall inflammation e.g., hypoxia, abnormal flow stresses and venous blood stasis as elicited by altered levels of nitric oxide and prostaglandins in circulation [48,49]. The mechanical status of the venous endothelium, its glycocalyx and the circulating blood cells are indeed critical for sensing changes ocurring in the venous bloodstream, and for the expression of membrane-receptor cytokines and other cell adhesion molecules necessary for normal venous circulation. In structural terms [50], the amphipathic nature of saponins outfits them with a potential ability to interact with membrane lipid components such as cholesterol, phospholipids and sphingolipids [39,51-53], which encode the functional molecular crosstalk among the structural fluid mosaicity that supports membrane activity [1-7].

**Figure 1.**
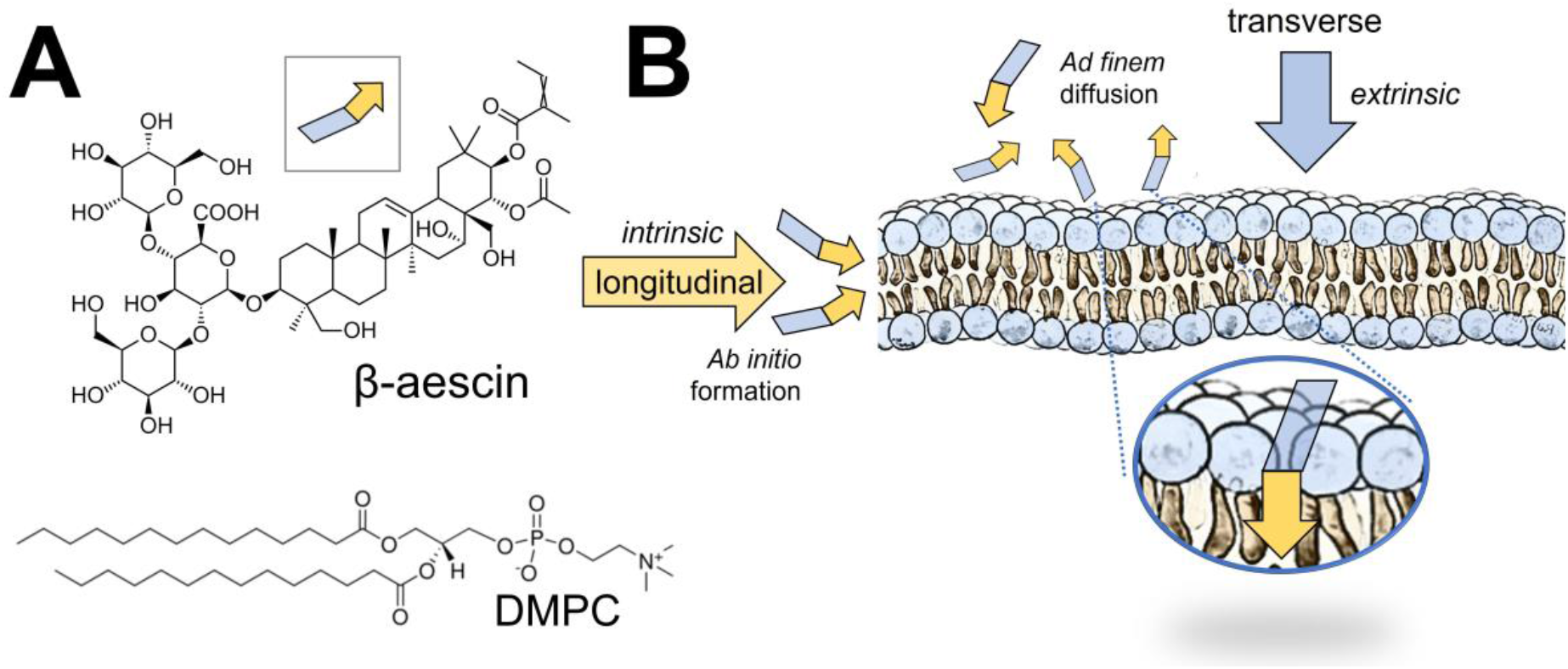
**A) Chemical formula** of β-aescin, or *escin* (M.W. 1,131.26 g mol^-1^). Notice the amphiphilic structure of the molecule with a polar head constituted by a trisaccharide sugar (blueish trapezoid), and a hydrophobic moiety of pentacyclic terpenoid (aglycone) basis prone to membrane insertion (yellowish arrow); see *inset*. DMPC is the structural phospholipid chosen to recapitulate a fluid and flexible membrane. **B) Scenarios of membrane insertion** proposed for *escin*. The intrinsic insertion corresponds to the *ab initio* integration of escin in the lipid formula before membrane formation (longitudinal escin insertion). The extrinsic insertion corresponds to the *ad finem* integration of escin by exogenous adsorption after membrane formation (transverse escin insertion). *Adsorption mechanism:* Although escin first interact with the phospholipid membrane through of its polar head, once adsorbed it reorganizes vectorially with the steroidal ring embedded in the hydrophobic core of the lipid membrane; see *inset* for a depiction for final equilibrium insertion. Both scenarios of escin-insertion are studied comparatively in this work.

Here, we demonstrate the genuinely mechanical influence of escin as lead by its incorporation in model systems based on phospholipid membranes, particularly Langmuir monolayers and bilayer vesicles as composed by the zwitterionic DMPC phospholipid. By exploiting DMPC in a minimal reductionistic approach to membrane mechanics, given its mesogenic transition between liquid-crystal bilayer phases near room temperature (from the ordered gel phase up to the disordered L_α_-phase at *T*_*m*_ ≈ 24 °*C*), their mosaic model membranes (cf. phase-separated monolayers and bilayers) both recapitulate structural resilience, fluidity and flexibility as being tunable in terms of experimental compositions and temperature. As depicted in Figure 1B, a modelling biomimetics essentially captures a simplified membrane in which escin can be inserted either as a “congenital” fabric within the intrinsic lipid formula (longitudinal insertion), or as an exogeneous adsorption from the external aqueous suspension (transverse insertion). Furthermore, the simplest Langmuir monolayers will be exploited as minimalistic membrane models for discriminating the spatiotemporal scales of mosaic organization in which escin operates dually either when extrinsically adsorbed from the aqueous phase or when intrinsically incorporated to the membrane. By using neutron and light scattering techniques with large unilamellar vesicles prepared by extrusion, we have previously shown how paradoxically escin causes mosaic liquid-solid domain segregation in rigid DMPC-bilayers leading to softer membranes by eliciting mesoscopic disorder under phase separation [39,51]. In the present study we are going beyond those indirect studies of escin impact, and by exploiting conventional techniques to track adsorption / insertion and surface rheology, together with membrane microscopy (BAM and fluorescence), we have provided experimental evidence on the existence of rigid domains mainly composed of escin. They trigger a dual membrane mechanics specifically controlled upon the escin insertion status. By considering the two escin penetration scenarios aforementioned, we have assessed the structural disruption of escin in DMPC bilayers constituted as giant unilamellar vesicles (GUVs). The membrane mechanics of the single lipid leaflets has been further explored in the corresponding Langmuir monolayers supported on an aqueous solution containing escin monomers. Our experimental results support the hypothesis on the (de)compacting incorporation of escin depending not only on the solid or fluid DMPC-phase state, but also on the vectorized direction particularly chosen for escin insertion. We have demonstrated that the pre-existing molecular ordering acts as a mechanical scaffold with a dual mechanical function i.e., a key regulator of membrane rheology thereby escin eventually triggers either a structural stiffening or a softening in dependence on the molecular orientation during the incorporation process.

## 2. Experimental section

### 2.1 Chemicals

The purified molecule β-D-Glucopyranosyl-(1→2)-[β-D-glucopyranosyl- (1→4)]-(22α-(acetyloxy)-16α,24,28-trihydroxy-21β-{[(2Z)-2-methylbut-2-enoyl]oxy}olean-12-en-3β-yl(β-D-glucopyranosidu-ronic acid), shortly named β-aescin (or simply escin) was purchased from Sigma (purity grade as a pharmaceutical primary standard), and used without further purification. The single escin molecules were dissolved in ultrapure water below the critical micellar concentration (CMC approx. 0.4mM). The fully saturated phospholipid 1,2-dimyristoyl-sn-glycero-3-phosphatidylcholine (DMPC), and the associated lipophilic dye 1,2-dioleoyl-sn-glycero-3-phosphoethanolamine-N-(lissamine-rhodamine B sulfonyl) (RhPE) were supplied by Avanti Polar Lipids. These lipids were dissolved in chloroform/methanol mixture to achieve a final concentration of either 0.1 mg/mL DMPC for experiments with Langmuir monolayers, or 1 mg/mL DMPC for preparing GUVs (0.1% by weight RhPE). Stock chloroform solutions were stored at -20 °C. All the chemicals were obtained for Merck-Sigma. Ultrapure water was obtained from a Milli-Q system (Millipore; resistivity higher than 18 MΩ cm; organic residual lower than 1 ppb; surface tension 72.6 mN/m at 20 °C). The experiments were performed in water solutions buffered with 50 mM phosphate (pH 7.0).

### 2.2 Escin monolayer penetration

To measure surface pressures in adsorption monolayers, we used a computer-controlled Langmuir trough (KSV/NIMA Small, 77.5 cm2, Biolin Scientific Holding Ab, Stockholm, Sweden). The Langmuir trough is equipped with a pressure measuring system with a paper Wilhelmy-type sensor and two mobile Teflon barriers. DMPC was deposited at the air/water interface using a chloroform spreading lipid solution (0.1 mg/mL). Prior to initiate measurements the monolayer was left to equilibrate by allowing the solvent to evaporate (approx. 15 min). The monolayer was then compressed to a nominal pressure *π*_0_. In extrinsic adsorption experiments, the subphase (50 mM phosphate buffer) was exchanged with a submicellar solution of escin (0.1 mM in phosphate buffer) using a peristaltic pump connected outside the barriers. The surface pressure versus time measurements were recorded continuously until a constant pressure was reached. All the experiments were carried out either at 4 ºC (low-T experiments), or at 38 ºC (high-T experiments).

### 2.3 Monolayer structure: Brewster angle and fluorescence microscopy

Brewster angle microscopy imaging (BAM) reveals the internal structure of the monolayer on the micrometric level. BAM measurements were performed in a Langmuir trough installed on a I-Elli2000 (Accurion GmbH) using a Nd:YAG diode laser (532 nm wavelength; 50 mW power). An incidence angle θ = 55.2 ± 0.1° was chosen in order to maximize the reflected intensity and the polarization sensibility. BAM images are taken simultaneously to the surface pressure monitorization. Image processing included a geometrical correction for spherical aberration, as well as a filtering to reduce interference fringes and noise. The reflected light in BAM imaging can be expressed by the Drude’s equation [54]:

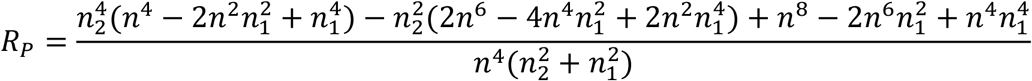

with *n* being the effective refractive index of the Langmuir monolayer film, *n*_1_ the refractive index of the air, and *n*_2_ the refractive index of the aqueous subphase in the near vicinity of the monolayer. Within this effective model not only the refractive index of the monolayer but also the amount of reflected light is proportional to the local thickness of the composite film supported on water. Consequently, the darker regions correspond to the locally thinner monolayers (or no presence of monolayer), whereas the brighter regions correspond to the thicker areas of the monolayer.

### 2.4 Isotropic compression isotherms: hydrostatic compressibility modulus

In order to measure the compressibility modulus under hydrostatic compression we exploited the conical cup device previously described in Ref. [10]. Briefly, the Langmuir monolayers were spread on the escin-containing subphase from a DMPC chloroform solution (1 mg/mL), and the solvent left to evaporate for 30 mins till monolayer equilibration. Once the spreading equilibrium was reached, the surface area available to the monolayer was progressively reduced by lifting up the conical cup whereas the surface pressure is measured at the center of the trough using a paper Wilhelmy plate attached to a force sensor (PS4, NIMA). This device allows the measurement of the changes in surface pressure under the isotropic compression produced by the radial contraction of the surface area available to the monolayer. The subphase temperature was controlled by recirculating water from a thermostatic bath (Polyscience) through a fluid circuit placed at the bottom of the trough and measured by a Pt-100 sensor. The experimental setup was placed in a transparent plexiglass box in order to avoid undesirable air streams and/or dust deposition on the surface during experiments.

### 2.5 Monolayer viscoelasticity: interfacial shear rheology

We performed measurements of two-dimensional shear rheology using a double-wall rheometer under thermostatic and evaporation control (DWR ARES-G2). To form the Langmuir lipid monolayer a small amount of DMPC dissolved in chloroform/methanol (1:1 v/v at 0.1 mg/mL) was spread as droplets onto the surface of the buffered subphase. To completely evaporate the chloroform an awaiting time of 30 minutes was used prior to the experiments. Then, a submicellar escin solution was injected into the subphase (0.1mM), and the system was left to equilibrate overnight. In order to avoid subphase evaporation, the Langmuir film was first tempered at 4ºC, then left overnight, and finally, the temperature adjusted. For low-T measurements, rheological Langmuir film experiments were performed at 4ºC. The stress in the samples was tracked permanently and the measurements at any temperature were only taken when the mean stress reached a constant value. Experimental replicas were performed in triplicate and the results analyzed corresponded to their arithmetic averages.

### 2.5 Giant unilamellar vesicles

The giant unilamellar vesicles (GUVs) were prepared by the electroformation method according to the optimized protocol described by Mathivet et al [55]. To prepare the GUVs, the lipid powder was first dissolved (at 1 mg/mL) in a chloroform/methanol mixture (2:1 v/v), then a drop of 20-30 uL was deposited on the ITO slide, and finally, the solvent was removed by evaporation in a stream of dry nitrogen. To prepare the GUVs of DMPC in the fluid state, we assembled ITO chambers first filled with an aqueous sucrose solution (200mM), then placed inside an oven at ca. 38°C, a temperature well above the melting temperature of the phospholipid (T_m_ = 23.6°C). Afterwards, the sealed electroformation chambers were connected to an electric field for three hours (8Hz, 1.8V). For subsequent GUV visualization under the optical microscope, the GUV sample previously electroformed in sucrose is further diluted in a glucose solution with a concentration slightly higher (208mM). Such density contrast favors properly both GUV sedimentation and visualization near to the microscopy slide. For the intrinsic insertion assays, methanol-dissolved escin was added (*ab initio*) to the lipid solution dissolved in chloroform. With respect to DMPC, we achieved a final escin concentration of 30 % w/w. In the extrinsic insertion assays, however, a solution of escin monomers was added (*ad finem*) once again the DMPC vesicles were formed. The final concentration of escin was reached in the suspending medium (40 µM). For fluorescence microscopy, a small amount of RhPE (0.1 % w/w) was added to the lipid solutions before preparing the lipid film.

## 3. Results

### 3.1 Longitudinal escin insertion elicits molecular ordering in fluid DMPC-based membranes

Escin is known to intrinsically leads to the formation of solid domains when incorporated *ab initio* into a fluid membrane formula [39,50,51]. It behaves as if it were integrated de novo through membrane biogenesis (intrinsic longitudinal insertion). To evaluate such “longitudinal” insertion into GUVs formed by a fluid DMPC membrane (at high temperature *T* = 38 °*C* > *T*_*m*_), we exploited the fluorescence emission of RhPE, the lipophilic fluorescent dye able to conjugate to fluid phospholipid phases of a disordered nature e.g., the liquid disordered phase [56]. Because the incorporation of escin triggers phase segregation into ordered arrangements [39,52], the fluorescent probe is excluded out from the *escin*-rich ordered phases e.g., liquid ordered (L_O_), gelly-like and crystalline solid phases. As assessed by confocal microscopy in the liquid disordered (Lα)-phase of the lipid bilayer (at *T* > *T*_*m*_), Figure 2A shows the formation of monophasic GUVs (1Φ) as revealed by the homogenous distribution of the RhPE dye (Fig. 2A; left panel). However, intrinsic *ab initio* incorporation to the binary GUVs were observed to partition escin and DMPC into a biphasic coexistence (2Φ), as discrete escin-rich solid domains (excluding out the fluorescent dye), dispersed in a continuous DMPC-rich fluid phase (as containing the RhPE dye). Only the fluid character of the liquid ordered (L_O_)-phase of the lipid bilayer composed of saturated hydrocarbon chains packed with local hexagonal order is expected to allow phase coexistence at equilibrium with the liquid disordered (Lα)-phase into spherical GUVS (right panel). Indeed, the bilayer Lα–L_O_ coexistence appears still fluidlike as far as the black domains appeared with the rounded boundaries continuing the liquid domains (see inset image). As intrinsically mixing components in a broad range of escin concentration (up to ca. 40% w/w), biphasic 2Φ-segregation was systematically observed in almost all GUVs present in a microscopy field (including encapsulated vesicles; see Fig. 2A, bottom). However, no stable GUVs were obtained at higher escin concentration (above 40% w/w), as expected for solidlike membrane with a high fragility insufficient to make the GUVs swelling. Therefore, our observations confirm the longitudinal integration of escin into DMPC to elicit a 1Φ→ 2Φfluid phase segregation into *escin-rich* L_O_-domains dispersed in the continuous *DMPC-rich* Lα-phase of the phospholipid bilayer (see schematic in Fig. 2A; bottom).

**Figure 2.**
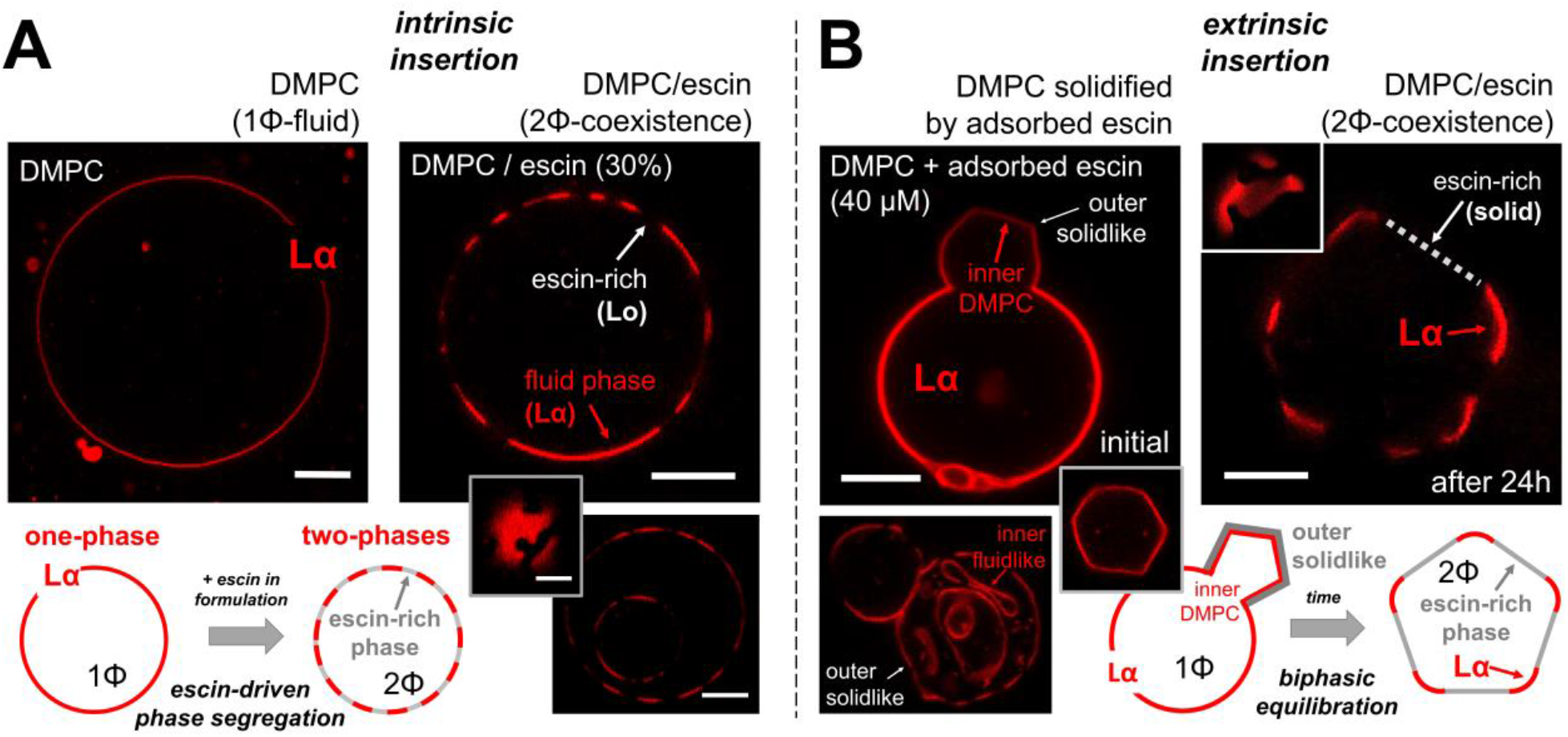
Fluorescence microscopy images obtained on the *escin*-insertion scenarios as designed in Fig. 1B for electroformed giant unilamellar vesicle (GUV) -membranes. The fluorescent dye RhPE is selective for the fluid disordered phospholipid Lα-phase. *Scale bars:* 2 microns. **A) Intrinsic *ab initio* insertion**, or longitudinal phase-separation scenario in which escin is included *ab initio* within the electroformation mixture. *Left panel)* A pure-DMPC vesicle in the homogeneous liquid disordered phase (Lα). *Right panels)* A hybrid, phase-separated (Lα-L_O_) vesicle as electroformed from the DMPC/escin mixture (30% w/w.). *Inset)* Zenithal view of a Lα-L_O_ separated vesicle showing circular L_O_-domains in a Lα-continuous. *Bottom)* Two “matrioska” vesicles appear with the same heterogeneous Lα-L_O_ configuration as formed *ab initio* from a phase-separating mixture. **B) Extrinsic *ad finem* insertion**, or transverse phase-separation scenario in which escin is membrane inserted *ad finem* by adsorption from the suspending medium. *Left panels)* Initial transformation stage thereby a pure-DMPC vesicle in the fluid Lα-phase becomes deformed under the action of a solid top layer locally adsorbed after 1 hour incubation (in 40 μM escin, below CMC). *Inset)* Polygonal edge profile of a solidified GUV under complete rigidization by an escin top layer. *Bottom)* Matrioska vesicles with only the outer membrane rigidized in contact with escin thus appearing solidlike (flat edges). The inner membranes remain fluidlike as they appear flexibly adapted to the internal space (rounded edges). *Right panel)* Final equilibrium of the DMPC/escin vesicles as Lα-L_O_ phase-separated (after 24h *escin-*incubation). Once adsorbed and further reorganized, the escin-rich L_O_-domains appear as flat edges excluding out the fluorescent dye. The DMPC-rich Lα-phase adapts the rigid flat domains as a continuous flexible junction (see inset). **Schematics**: The bottom panels systematize the observed behavior in each insertion scenario.

### 3.2 Transverse escin insertion induces extrinsic membrane stiffening following solidlike decoration in DMPC-based bilayer vesicles

We are now interested on how escin impacts phase coexistence when incorporated *ad finem* in previously formed DMPC vesicles –as if escin was extrinsically inserted from the outer medium (transverse insertion). Figure 2B shows the rationale designed for this purpose. We first prepared pure-DMPC GUVs at the liquid-disordered L_α_-phase (without escin included in the lipid formula). Then, we added escin monomers dissolved sub-micellar in the vesicle outside (at 40μM final concentration). Since no escin micelles were present to disentangle the lipids from the GUV bilayers (at *c* ≈ 0.1*mM* ≪ *CMC*), the hybrid DMPC / escin GUVs remain completely stable in escin suspension. At adsorption interaction with the outer escin monomers, the hybrid 1Φ-GUVs initially remained in the disordered L_α_-phase –as far no phase segregation is observed during the first stages of escin adsorption (Fig. 2B; left panel). However, at longer times within the first hour after escin adding, we observed most GUVs losing circularity (e.g., polygonal profile in the inset image). The *escin*-modified DMPC-GUVs progressively formed flat edges that would suggest a solid character of the outer adsorbed escin layer, although maintaining a disordered lipid arrangement in the DMPC bilayer underneath (see Fig. 2B; left). Because the escin monomers primarily adsorb on the outer vesicle surface they can freeze *in situ* as solid planar plates [40], thus forcing the underlying DMPC membrane to adapt its extrinsic geometry without changing the intrinsically fluid composition than enables its natural roundness (Fig. 2B; left). As additional evidence for this extrinsic interaction during the initial adsorption stage, where the DMPC bilayers stay protected from escin insertion (akin “matrioska” vesicles), they remain fluidlike with a rounded profile without flat edges (Fig. 2B; bottom). However, when allowed to reach adsorption equilibrium (after incubation for 24h with escin), these vesicles undergo biphasic 2Φ-segregation now observed as an exclusion of the fluorescent probe from the blacked (escin-rich) areas where the flatter geometries suggest a solidlike character –the planar edges. However, the colored (DMPC-rich) areas remain fluidlike as corresponding to the presence of fluorescent dye in the disordered Lα-phase –the rounded vertices (see Fig. 2B; right panels). These observations reveal how the transverse *escin*-integration in DMPC membranes follows the same two-stage adsorption / reorganization mechanism as observed in other fluid membranes [28,57,58]. Firstly, escin-adsorption induces extrinsic membrane stiffening in the flexible (Lα) DMPC-membrane in which it is becoming adsorbed *–primary relaxation*. Secondly, once adsorbed escin has enough time to reorganize and equilibrate with the DMPC-bilayer, the hybrid membrane so formed then undergoes phase segregation from 1Φ(Lα-phase), up to 2Φ(Lα-L_O_ phase demixing) *–secondary relaxation*. Under final equilibration, the rigid L_O_-domains appear solidlike into the flattened vesicle edges becoming mechanically equilibrated through of flexible joints constituted by rounded vertices of the continuous L_α_-phase (see schematic in Fig. 2B; bottom). The L_o_-domains are presumably *escin*-rich as far the fluorescent dye is excluded out from the laterally ordered hydrophobic arrangement constituted as π–π stacks between the steroidal moieties of the β-aescin molecule [39,59]. In agreement with the previous physicochemical hypotheses [39,50,60], our results suggest the equilibrium state of escin inside the phospholipid bilayer as being reached through the direct interaction between the acyl chain hydrophobic part of the membrane and the aglycone counterpart of the rigidizing molecule akin the structurally reinforcing interaction between phospholipids and sphingolipids with cholesterol in real biomembranes [4,7].

### 3.3 Transverse escin penetration induce crystallization in DMPC monolayers

To confirm the solidlike crystalline nature of the membrane escin-rich domains at low temperature (*T* = 4°*C* ≪ *T*_*m*_), we studied the transverse incorporation of *escin* in a simplified crystallization model based on Langmuir DMPC monolayers penetrated from below by the *escin* monomers dissolved in the subphase (see Methods). The fluorescence images in Figure 3A evidence the formation of black polygonal domains (excluding the dye outside), which evolve in time as a nucleation process for crystal formation as growing from the continuous fluid phase. Over the days passed after monolayer formation, we observed how these solid domains further grow and mature as polygonal plates with very sharp edges (see Fig. 3A; final zoom image of *escin*-crystals). The sequence observed as a crystal nucleation and growth process suggests a driving mechanism of surface reorganization of the monomers diffusing from the bulk solution although somewhat controlled by some kind of adsorption energetics (Fig. 3B). Such regulatory dynamics could be dependent either on the availability of adsorption sites in the preformed DMPC / escin monolayers (entropic mechanism), or on their capacity to attract cohesively escin (enthalpic mechanism). We performed a control experiment to ensure that *escin* incorporation as crystals into the DMPC monolayer is not due to the presence of RhPE, whose small lipid head could arguably induce DMPC or *escin* to crystalize as a nucleation seed. An additional experiment confirming no crystal formation was performed at high temperature (*T* = 38°*C* ≫ *T*_*m*_); no membrane domains of any kind are observed in high-T states of the hybrid DMPC / escin monolayers. These negative controls have completely ruled out the possibility for structural artifacts that could complicate the interpretation of our dynamic results in adsorption monolayers. As a further insight going beyond the above qualitative analysis on low temperature crystallization, we studied the transversally organized incorporation of escin by taking advantage of the dye-free visualization enabled under the Brewster angle microscope (BAM). After DMPC monolayer formation in a homogenous L_O_-state at low temperature (*T* = 4°*C* ≪ *T*_*m*_), escin was further injected into the subphase under identical conditions as above leading to the results in Fig. 3A (except for the absence of dye). After 12h incubation, the small solid domains started to nucleate (Fig. 3C; 12h panel), with an irregular sharp-edged texture typical for nucleating crystals (see *inset*), and a size similar to that observed in the presence of dye (see Fig. 3A; panel for 12h). As regards the BAM-contrast between the solid crystals and the liquid surrounding, we always detect the solid domains with an optical aspect significantly darker than the continuous liquid phase (Fig. 3C; inset). The average ratio of BAM intensities was measured at *I*_*liquid*_/*I*_*solid*_ = 1.4 ± 0.2, indicating a monolayer thickness smaller in the solid crystals than in the liquid phase (see Methods). These results essentially indicate a fluid phospholipid monolayer thicker in head-to-tail length than the dispersed crystals assumed merely composed by the thinner *escin* molecule (see schematics in Fig. 3C). Regarding specific molecular dimensions in monolayer arrangements, we expect a stretched molecular length for DMPC, 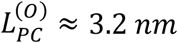 (including the hydrated polar head as corresponding to the monolayer L_O_-phase equivalent to the bilayer gel phase [4,5]); and a shorter length for escin 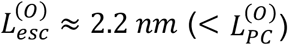 (see Fig. 3C; right panel). As aforementioned, Figure 3C depicts these differences in molecular sizes as inferred from molecular models which generalize structural data for escin adsorption monolayers using neutron scattering and x-ray diffraction [52,53]. From this molecular modelling we indeed estimate the thickness ratio 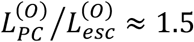 which is compatible with the measurement of BAM contrast between the two monolayer phases observed at coexistence (liquid ordered DMPC-rich (L_O_), and solid crystalline escin-rich); this is given indeed by the ratio of monolayer thicknesses i.e., *I*_*LO*_/*I*_*solid*_ = 1.4 ± 0.2 ≅ *L*_*PC*_/*L*_*esc*_ ≈ 1.5.

**Figure 3.**
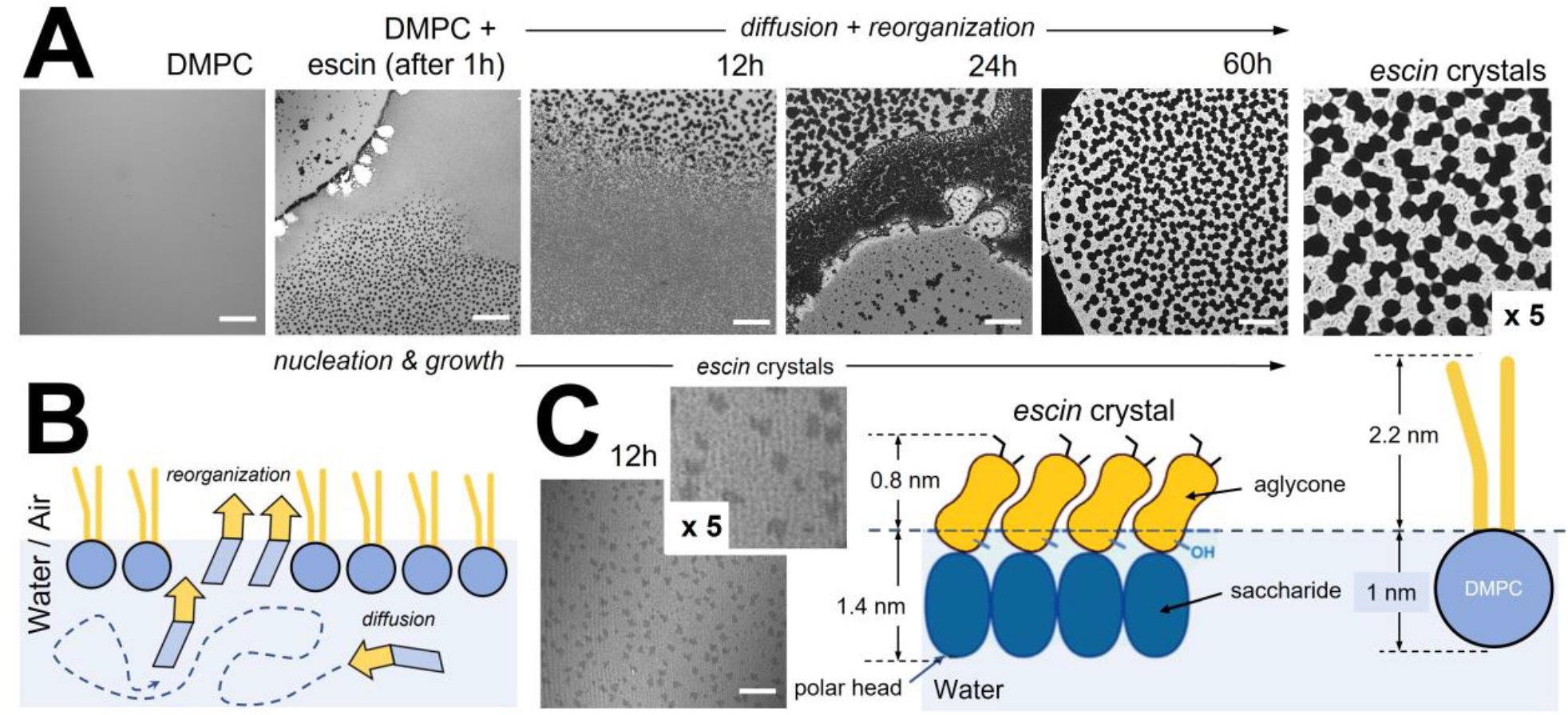
**A) Crystal nucleation and growth process at low temperature**. The sequence shows the time evolution of the penetrating interaction of *escin* with a disordered DMPC-monolayer fluorescently labelled with RhPE 0.1% mol (in fluorescence microscopy the DMPC-richer, the brighter; *Scale bars:* 20 microns). The first microscopy snapshot corresponds to the bare DMPC monolayer. Then, escin is injected in the subphase and the fluorescence images captured at different times along 3 days. A nucleation-growth process is recorded as small grains of ordered phase (*escin*-rich; black) appearing larger and darker with longer times over an increasingly brighter background corresponding to the disorder phase (DMPC-rich; white). *Right zoom)* After 60h escin adsorption, the hybrid DMPC-escin monolayer reaches a steady appearance as escin-crystals with a homogeneous morphology in equilibrium with the DMPC liquid disordered phase (x5 zoom). **B)** Schematics for possible escin adsorption scenario from a water subphase to a lipidic Water/Air (W/A-) interface. The soluble surfactant molecule (escin) diffuses from the bulk to the W/A-interface. Here, it penetrates and reorganizes to reach the equilibrium with the lipid component. **C)** Brewster angle microscopy (BAM) of the DMPC monolayer penetrated by escin 12h after injection in the subphase. The nucleation of monolayer *escin-*crystals is evidenced with a thickness smaller (darker) than the DMPC-continuous phase (brighter). *Inset)* The nascent crystals are quite heterogeneous in size (0.2 micron in average), at compatibility with the fluorescence images at the same adsorption time (bottom image in A). **Molecular model**. The *escin-*crystals are formed by hydrophobic stacking between the core rings in the aglycone moieties (van der Waals), and hydrogen bonding between the saccharide heads dangling in water. This molecular modelling assumes a locally thinner monolayer at the level of the *escin-*crystals than found in the thicker disordered phase mainly composed by DMPC in the ordered phase (see molecular dimensions).

### 3.4 Escin membrane insertion depends on the DMPC-monolayer packing status as determined by surface pressure and temperature

The above observations on escin-crystal formation in a biphasic system at low temperature suggested a two-step mechanism mediated by an initial nucleation and the further crystalline growth as driven by a bulk-to-surface diffusive adsorption kinetics limited by a reorganizational energetics that determines the final equilibrium (see depiction in Fig. 3B). Although a similar adsorption kinetics certainly happens at higher temperatures, the surface system remains monophasic as a disordered fluid phase. In a previous paper we indeed evidenced a preference of escin for heterogenous demixing in DMPC monolayers at the lower temperatures studied [39]. Here, we have explored this behavior in detail and observed an inducing role of ordered phospholipids for *escin*-insertion into quasi-2D crystaline monolayers. We have studied the kinetics of adsorption at the water/air (W/A)-interface as a function of the packing state of the lipids on the supporting DMPC membrane. Figure 4A shows adsorption plots recorded at both temperature regimes: *Low-T)* at a temperature of 4ºC, at which the phospholipids chains become tightly packed as mesogenically arranged in an ordered state (L_O_-phase); *High-T)* at 38°C, in which DMPC chains are in a disordered state (L_D_-phase). By considering these experimental kinetic plots, the diffusion of escin is revealed initially slower at lower pressures whilst the final equilibrium for membrane escin-incorporation is reached at higher pressures. This kinetic bimodality effect becomes even more pronounced when both the lipid packing, and temperature is lower (Fig. 4A; left panel). In Figure 4B, this difference in incorporation is the best evidenced when we analyze the equilibrium insertion pressure as a function of initial pressure (*π*_*ins*_ *vs. π*_0_ plots). Here, we evidence that when the lipids are arranged in an expanded L_D_-phase, there is hardly any significant difference in the escin incorporation between low and high temperatures. However, when the initial pressure is higher than 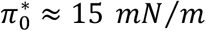 (as corresponding to the L_D_ → L_O_ transition), the higher equilibrium insertion pressure happens in the L_O_-phase at lower temperature. Arguably, this effect is due to a reduced *escin* mobility in the compacted phase allowing an enhanced steric interaction.

**Figure 4.**
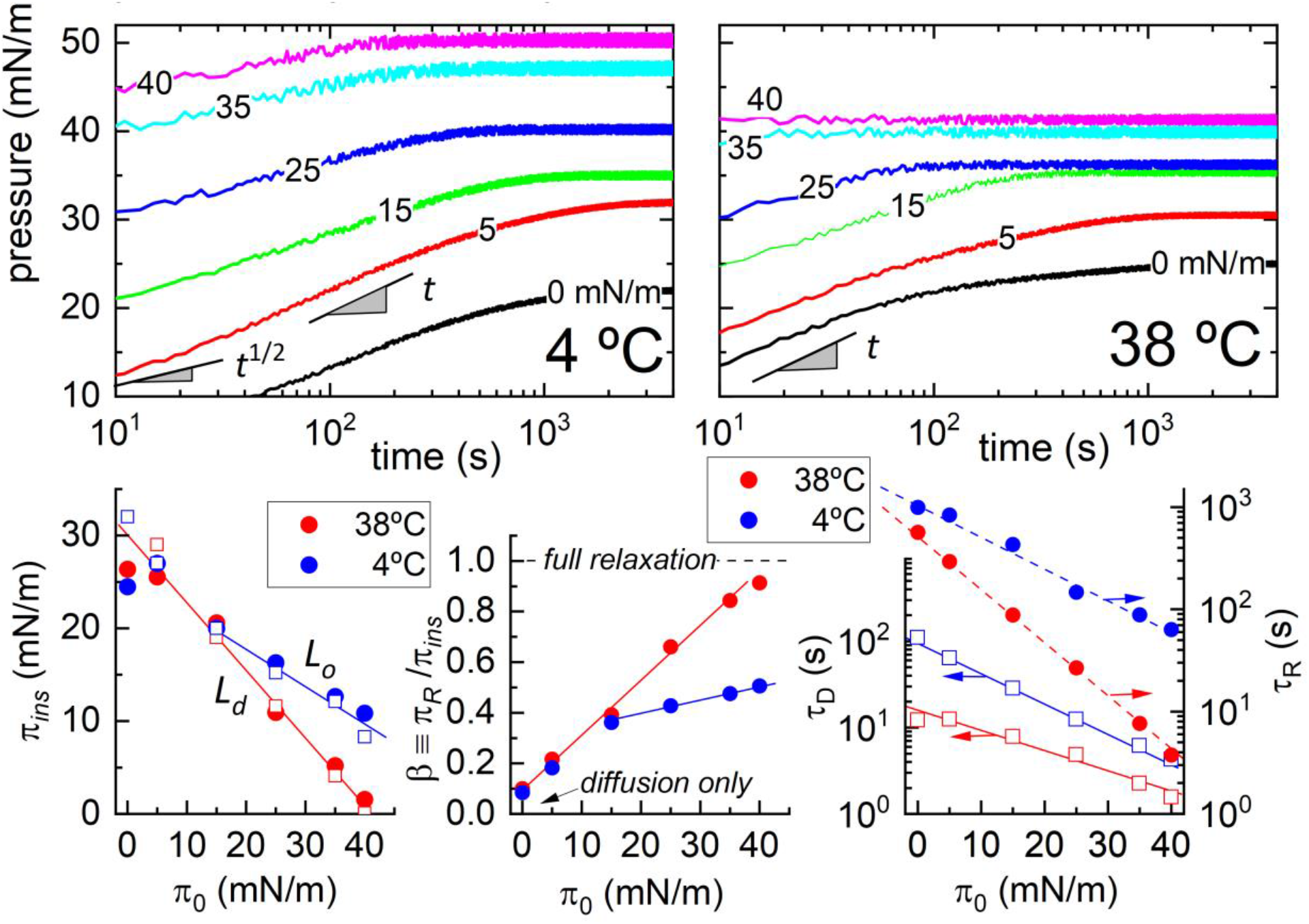
**A) Adsorption kinetics** of *escin* on a DMPC monolayer at different packing states (Π_0_ = 0, 5, 15, 25 and 30 mN/m). Escin is dissolved at constant bulk concentration (ca. 0.1 mM), and the system considered at 4ºC (left) and at 38ºC (right). Best fitting parameters obtained from Π(t) adsorption kinetics (see Eq. 1). **B) Equilibrium insertion pressure** Π_ins_ as a function of initial pressure Π_o_ upon *escin* incorporation into the DMPC monolayer. **C) Energy balance** between the amplitude of internal reorganization and the insertion pressure at equilibrium as a function of initial pressure Π_o_. **D) Adsorption kinetics parameters:** diffusion times (left axis) and membrane reorganization times (right axis) as a function of temperature and initial pressure Π_o_.

### 3.5 Escin penetration kinetics: adsorption follows membrane reorganization

According to the adsorption theory by Graham and Phillips (GP), the study of the penetrating relaxation pressures for given initial states provides information about the characteristic times associated with the adsorption and subsequent incorporation of penetrating escin surfactant in the W/A-interface [61]. As commented above, Figure 4 showed the experimental results on the adsorption kinetic plots *π*(*t*), as tracked for different penetration status of escin in the DMPC monolayer considered at variable initial pressure (*π*_0_), and temperature (*T*), for fixed sub-micellar escin concentration (*c* = 0.1 *mM* ≈ 0.25 *CMC*) [40]. The penetration mechanism followed an obvious two-step kinetics i.e., an adsorption–reorganization process thereby the pressure jump with respect to the initial status of the adsorbent surface is given as *π*(*t*; *π*_0_) = *π*_0_ + *π*_*P*_(*t*; *π*_0_, *c*) [57]; under penetrating relaxation, the GP-kinetic equation follows [61]:

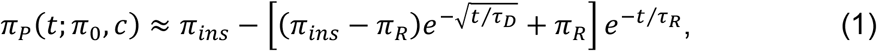

being *π*_*ins*_(*π*_0_, *c*) the equilibrium insertion pressure of escin, and *π*_*R*_(*π*_0_) ≤ *π*_*ins*_(*c, π*_0_) the partial amount of escin-stabilizing surface energy as due to its internal reorganization.

The two relevant GP-processes do appear consecutively, first, an initial adsorptive transport characterized by a diffusive time *τ*_*D*_(*c, π*_0_), which is followed by a subsequent terminal relaxation under *escin-*penetration at longer reorganization times *τ*_*R*_(*π*_*ins*_) ≫ *τ*_*D*_(*c, π*_0_). Because the dynamical changes in surface pressure occurs under energy conservation a further balance equation can be written as:

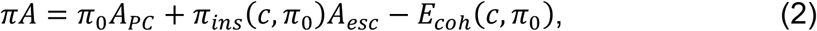

being −*E*_*coh*_ = *π*_0_(*A*(*c*) − *A*_*PC*_) a cohesion energy experienced by the surface molecules occupying an average area *A* under escin insertion; for *A* = *A*_*PC*_, then *E*_*coh*_ = 0.

Upon thermodynamic equilibrium *Δ*(*πA*) = 0, one expects fulfill a cohesive energy trade-off as due to escin and DMPC interacting each other under initial pressure (*π*_0_):

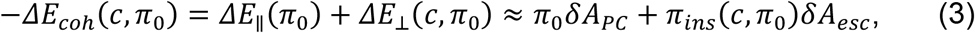

as due to compacting changes in the respective surface areas *δA*(*π*_0_) = *A*^(*init*)^(*π*_0_) − *A*^(*final*)^(*π*) (> 0). All the above constitutive parameters depend on the initial pressure (*π*_0_), which makes the insertion pressure to decrease conversely i.e., *π*_*ins*_ = *π*^(0)^ − *απ*_0_. This effectively linear relationship is considered with a positive intercept due to cohesive energy by adsorbed escin (*π*^(0)^ ≡ −*ΔE*_*coh*_/*δA*_*esc*_ ≫ 0), and a near unity slope under relative area compaction (*α* ≡ *δA*_*PC*_/*δA*_*esc*_ ≈ 1).

These constitutive relationships imply, on the one hand, a bulk-to-surface diffusivity controlled by an adsorption potential well of cohesive energy susceptible to declining with decreasing the availability of surface free sites for escin cohesivity (i.e., *E*_*ads*_ → 0 for *A* → *A*_*PC*_ at *π*_*ins*_ → 0). In other words, the adsorption energy becomes effectively attractive under transverse insertion of escin. Such an out-of-plane cohesive interaction in an attractive potential well results similar to the adhesive case considered by the BET theory of adsorption [62]. Surface adhesion due to inserted escin facilitates adsorption from the bulk phase into the DMPC-based surface as *E*_*ads*_ ≈ *π*_*ins*_(*A*_*PC*_ − *A*) ≪ 0 ⇒ −*ΔE*_⊥_(*π*_0_) ≈ −*απ*_0_*δA*_*esc*_ (for *π*_*ins*_ ≫ *π*_0_ and *δA*_*esc*_ > 0). Consequently, the initial adsorption times become effectively accelerated as due to a cohesively improved escin insertion i.e.,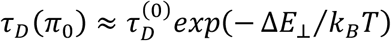 [62], upon transverse adhesive attraction leading escin compaction (under *δA*_*esc*_ > 0, this is Δ*E*_⊥_ = *π*_0_*δA*_*esc*_ ≫ 0, for *π*_0_ ≪ *π*_*ins*_). On the other hand, the terminal relaxation is expected as a lateral membrane reorganization of inserted escin which further becomes longitudinally reorganized. Consequently, an in-plane compaction energy globally favors the insertion of escin at approximate dependence on the pre-existing DMPC longitudinal packing i.e., −Δ*E*_*coh*_ ⇒ Δ*E*_∥_ ≈ *π*_0_*δA*_*PC*_ (for *π*_0_ ≫ *π*_*ins*_ → 0). Hence, a first-order reorganization kinetics appears within the DMPC-supporting structure under Debye-like relaxation times 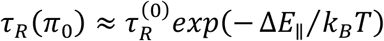 also upon cohesive attraction effectively leading more compact phospholipids (under *δA*_*PC*_ > 0, this is Δ*E*_∥_ = *π*_0_*δA*_*PC*_ ≫ 0 for *π*_0_ ≫ *π*_*ins*_). This longitudinal reorganization process becomes evidently faster with increasing lateral compaction (increasing *π*_0_), which enables lower insertion pressures (decreasing *π*_*ins*_). The pressure trade-off imposed by equilibrium leads to higher membrane rigidness under lower initial ordering by DMPC, but higher ordering scaffolded by inserted escin (see Eq. 2). Hence, we expect the surface stiffness varying under uniaxial deformations by the respective molecular compactions *δA*_*PC*_, and *δA*_*esc*_ [10,28,63]; as coupled in series (*δA* = *δA*_*PC*_ + *δA*_*esc*_), the surface rigidness per molecule stems on the total surface interaction as *εδA* ⇐ *Aδπ*, being *ε* an effective bulk modulus for membrane dilatation [64]; see Section 3.6). Hence, the effective adsorption rate is globally evaluated as an in-series sequence of insertion and reorganization events i.e., 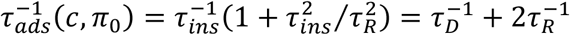 Therefore, two relaxation regimes can be experimentally inferred from the adsorption kinetics observed in limiting cases:

1. *Diffusion controlled* at short insertion times (i.e., *τ*_*D*_ ≈ *τ*_*ins*_ ≪ *τ*_*R*_, thus *τ*_*ads*_ ≈ *τ*_*D*_). In this slow relaxation regime (*τ*_*R*_ ≫ *τ*_*D*_), the initially quick insertion time is controlled by the faster diffusional adsorption (i.e., *τ*_*ins*_ ≈ *τ*_*D*_) [65,66], which becomes accelerated by the potential well of adhesion energy (−*E*_*ads*_ ≫ 0). Hence, the penetration pressure is dominated by the insertion pressure as 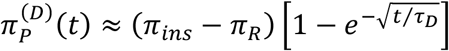 (for *π*_*ins*_ ≫ *π*_*R*_). This diffusion-controlled regime is clearly discerned scaling as 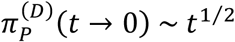 (see Fig. 4A; left panel), corresponding to low temperature and low initial pressure allowing for transverse escin-insertion (for *π*_0_ ≪ *π*_*ins*_; Fig. 4B).
2. *Insertion controlled* under internal reorganization (i.e., *τ*_*R*_ ≪ *τ*_*D*_ ≈ *τ*_*ins*_, thus *τ*_*ads*_ ≈ *τ*_*R*_/2). In this terminal relaxation regime (*τ*_*R*_ ≪ *τ*_*D*_), the penetration pressures increase as 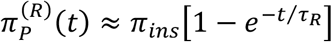(for *π*_*ins*_ ≈ *π*_*R*_) as controlled by the global cohesive energy that maintains the membrane internally organized (−*E*_*ads*_ ≪ −*E*_*coh*_ ≪ 0). The terminal insertion follows a first-order reorganization kinetics with an insertion time nearly twice the internal reorganization time (*τ*_*ins*_ ≈ 2*τ*_*R*_). The adsorption kinetics within this reorganization-controlled regime varies linearly on time as 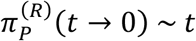 (Fig. 4A; right panel), being predominant at high temperatures and the relatively high pressures dominated by a longitudinal escin-reorganization (for *π*_0_ ≫ *π*_*ins*_; Fig. 4B).

The kinetic results shown in Fig. 4 have evidenced the two sequential regimes predicted by the GP-theory and somewhat inferred from the qualitative analyses of the microscopic membrane textures (see Figs. 2-3): *i)* the initial adsorption of escin towards the extrinsic bulk vicinity of the adsorbing surface; *ii)* the subsequent intrinsic incorporation of escin to the phospholipid layers. Below, we further analyze the corresponding energetics and relaxation times as determined by the initial pressure of the monolayer model in differential statuses of DMPC compaction as determined by temperature.

#### Adsorption energetics

The equilibrium pressures in Fig. 4B evidence the energy trade-off predicted between final escin insertion and initial DMPC state (see Eq. 2); for constant escin concentration, we detect in both phases the linear relationship 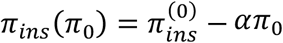 The positive intercept indicates a cohesive dominance of adsorption by inserted escin 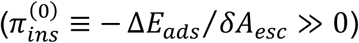 whereas the negative slope indicates higher escin compaction as referred to the saturated DMPC phospholipid (*α* ≡ *δA*_*PC*_/*δA*_*esc*_ < 1 i.e., *δA*_*esc*_ > *δA*_*PC*_). The best-fitted values obtained for these parameters from the measured data are collected in Table I, in reference to the different *L*_*D*_ / *L*_*O*_ states of structural compaction.

**Table I.** Energetic surface compaction parameters for the DMPC/escin system (as determined from the experimental data in Fig. 3B). The structural parameters for the corresponding cross-sectional areas are taken from literature data (as in Section 3.3): for DMPC [4,5]; for escin [39,42].

| **Surface phase** | $\pi_{ins}^{(0)}$ (mN/m) | $\alpha \equiv \delta A_{PC}/\delta A_{esc}$ | $A_{PC}$ (nm<sup>2</sup>) | $A_{esc}$ (nm<sup>2</sup>) | $-\Delta E_{ads}^{(esc)} \approx \pi_{ins}^{(0)}\delta A_{esc}$ |
| --- | --- | --- | --- | --- | --- |
| *Liquid disordered* | $30 \pm 1$ | $0.75 \pm 0.12$ | $\approx 1.0$ | $\approx 4.4$ | $35k_B T$ |
| *Liquid ordered* | $24 \pm 2$ | $0.56 \pm 0.15$ | $\approx 0.5$ | $\approx 0.9$ | $28k_B T$ |

At structural compatibility with the high- / low-temperature states of DMPC (L_D_ / L_O_), we detect clear differences rather controlled by the higher molecular malleability of reconfigurable escin (see schematics in Fig. 3), whose compaction changes are expectedly larger in the fluidlike L_D_-phase than in the solidlike L_O_-phase (see Ref. [53] for a detailed discussion); we found, respectively, for initial pressures 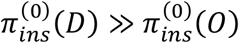 and for compaction slopes *α*_*D*_ > *α*_*O*_. By assuming structural data in Table I, we have estimated *δA*_*PC*_ ≈ 0.5 *nm*^2^, and *δA*_*esc*_ ≈ 3.6 *nm*^2^ ≫ *δA*_*PC*_, in qualitative agreement with the values found for the compaction ratio (see data for *α* in Table I). We have also estimated the escin cohesive interaction energies *i*.*e*., *the depth of the adhesive potential well for escin*, either relatively low in the ordered L_O_-phase at low temperature 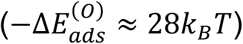 or a little higher in the L_D_-phase at high temperature 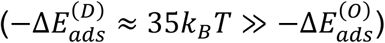 The observed differences reveal a higher escin-DMPC cohesion in the (L_D_)-phase at high temperature than in the (L_O_)-phase at low temperature (see Fig. 4B), which evidence the more facilitated escin insertion at conditions of higher surface disorder and / or high temperatures (higher adsorption availability for escin). Nevertheless, the kinetic data in Fig. 4C also evidence a difference in the relaxation amplitudes for escin reorganization *β*(*π*_0_) ≡ *π*_*R*_/*π*_*ins*_; although increasing also linearly with the initial pressures in the DMPC monolayers (*π*_0_), they experience a clear breakdown as spanning across the L_D_ → L_O_ transition. As above explained accordingly to the GP-adsorption dynamics, the values measured for *β*(*π*_0_) estimate how much the subsidiary reorganization step does compare to the primary diffusive transport at dependence of the membrane coverage status (see Eq. 1). At low surface coverage, escin insertion is chiefly dominated by external diffusion (i.e., at *π*_0_ ≈ 0, then 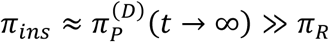 thus *β* ≈ 0). In highly covered surfaces, however, escin insertion mostly dominated under internal reorganization (i.e., at 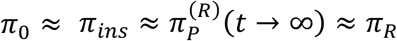 thus *β* ≈ 1). As clearly revealed by the kinetic plots in Figs. 4A (and the corresponding relaxation amplitudes in Fig. 4C), the process is limiting free-diffusive only upon conditions for practical absence of lipid at the interface (at *π*_0_ → 0). However, a progressive reorganization happens independently of the initial pressure in the expanded disordered status (at 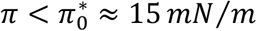). A tipping point is detected nearly the L_D_ → L_O_ transition as the onset of ordering for enhanced escin compaction in the L_O_-phase of the phospholipid (ca. 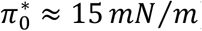). At that tipping point, the phase coexistence emerges as the lipid concentration increases upwards (at 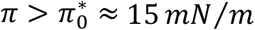 for *T* ≤ *T*_*m*_ ≈ 23°*C*). In these biphasic L_O_ / L_D_ states, the second reorganizational process appears to become the most important in the lipid-covered surface. At low temperatures (*T* ≪ *T*_*m*_), the organization of escin into the membrane is hence controlled either by the extrinsic diffusivity of the escin molecules towards the interface plane (in the L_D_-phase at low insertion pressures 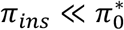), or by their intrinsic reorganization inside the membrane itself (in the L_O_-phase at high insertion pressures 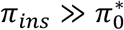). However, at the high temperature states (*T* ≫ *T*_*m*_) *–the most relevant from a biological point of view*, the subsidiary process for escin-insertion leads kinetics as a dominantly intrinsic reorganization. Henceforth, we consider the biologically relevant state to be closely corresponding to the liquid-crystalline ordering of the typical fluid lipid bilayer at nearly physiological temperature [5-7]. As referred to the ordered molecular packing in DMPC monolayers (L_O_-phase), this ideal reference state would correspond to high temperature (*T*_*high*_ = 38°*C* ≫ *T*_*m*_), and high pressure characteristic for the bilayer-like status 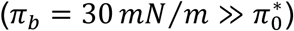 approximately half the maximal surface pressure reachable by a Langmuir monolayer before collapse [7]. These biomimetic conditions correspond to an averaged surface area *A*_*PC*_ ⇒ *A*(*L*_*O*_) ≈ 0.5 *nm*^2^ (see Table I), close to the molecular cross-section of a single phospholipid molecule in the liquid ordered (L_O_)-phase [5-7].

#### Adsorption kinetics

The kinetic data in Fig. 4D shows the dependency of the escin-specific relaxation times as experimentally derived from Eq. 1, respectively, *τ*_*D*_(*π*_0_) for the initially faster membrane-extrinsic diffusion (left axis), and *τ*_*R*_(*π*_0_) for the subsidiarily slower membrane-intrinsic rearrangements (right axis). The experimental values of the characteristic times and associated energies are collected in Table I, in reference to the different *L*_*D*_ / *L*_*O*_ states of structural compaction.

We found the characteristic relaxations becoming faster with increasing temperature (*T*), and with the initial pressure imposed by DMPC in the monolayer membrane (*π*_0_). At the low temperature (*T* = 4° *C* ≪ *T*_*m*_), the two adsorption times, either for the initial surface diffusivity or the subsequent reorganization, decrease exponentially (with increasing *π*_0_); indicating a higher surface adhesivity with increasing membrane compaction (Fig. 4D; blue symbols). At the high temperature (*T* = 38 °*C* ≫ *T*_*m*_), the two characteristic escin-insertion times decrease exponentially as well, however, as evidenced by a higher decaying slope, the reorganization time displays a marked tendency to further decreasing at the higher temperature (Fig. 4D; red symbols). Because the terminal reorganization is kinetically governed by the in-plane relaxation transport of DMPC and escin i.e.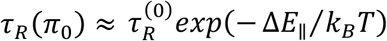, under lateral interaction energy −Δ*E* ⇒ −Δ*E* ≈ *π*_0_ *δA*_*PC*_, we hence expect the faster reorganization as due to the much stronger compaction. This high temperature behavior appears induced by escin at relatively low insertion pressure on the highly disordered (and very expanded) L_D_-phase of DMPC (i.e., 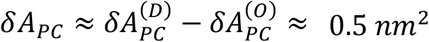 for 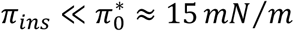). As comparing the GP-theory with the reorganization times experimentally observed (see Fig. 4D), we found the longest ones for the lowest temperature corresponding to the most escin-inserted states (escin rich L_O_-phase; i.e., *τ*_*R*_(*O*) ≫ *τ*_*R*_(*D*)), as corresponding for the most cohesive compactions induced by escin on the disordered phase of DMPC. As roughly estimated for the extrapolated reference state of no escin insertion (*π*_*ins*_ → 0 *mN*/*m*), corresponding to the bare DMPC molecules in the closest-packed configuration (at *π*_0_ → 40 *mN*/*m*), the lateral compaction parameter is experimentally determined in agreement with structural expectations i.e., 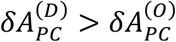 The energy for lateral reorganization is estimated higher in the more malleable disordered phase 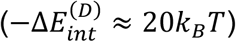 than in the liquid ordered status in which it appear as a cohesive energy for in-plane compaction (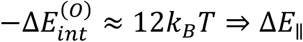; see Table II).

**Table II.**
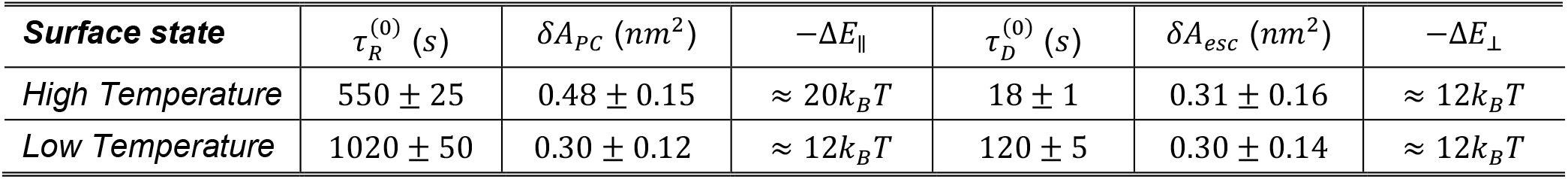
Experimental kinetic parameters for the DMPC/escin system in the high- / low-temperature states (as determined from the experimental data in Fig. 3D). Estimated interaction energies are per molecule.

Despite the longitudinal reorganizational differences upon escin insertion, the faster diffusional times showed differences dictated by the transverse transport from the subphase towards the adhesive surface; this is 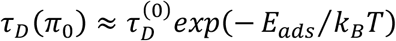 under effectively adhesive adsorption energy (i.e., *E*_*ads*_ < 0) [62]. This adhesion activated adsorption is essentially determined by the capacity of escin to create area when cohesively adhered to the interface i.e., Δ*E*_⊥_ ≈ −*π*_0_*δA*_*esc*_ (for *δA*_*esc*_ > 0). In experimental terms, we observed nearly the same slope independently of temperature *δA*_*esc*_ = 0.3 ± 0.1*nm*^2^ (see Fig. 4D; left axis), which is compatible with the compaction parameter measured for DMPC in the liquid order state. By assuming the maximal structural compaction above estimated for the membrane components i.e., *δA*_*esc*_ ≈ *δA*_*PC*_ ≈ 0.3 *nm*^2^, we estimate the transverse interaction parameter as an out-of-plane cohesive energy on the same order than the lateral cohesion energy; for the closest-packed state (*π*_0_ = 40 *mN*/*m*), these are Δ*E*_⊥_ ≈ Δ*E*_∥_ ≈ 12 *k*_*B*_*T*. This neatly attractive behavior evidences the surface adsorption of escin as an activated process of surface adhesivity essentially governed by the adsorption-enhanced diffusive transport from the bulk solution towards the adhesive surface. As being a matter of theoretical conjecture inferred from the above experimental observations, the amphipathic molecule β-escin should appear being highly surface active and self-adhesive as promoting high membrane cohesion due to lateral π-π interactions largely enhanced with increasing packing (*π*_0_).

### 3.6 Mechanical impact of escin-rigidity in Langmuir lipid monolayers

The solid-like behavior of the bare β-escin molecule has been well-established in previous studies in Langmuir monolayers and bilayer vesicles of the pure component [40,67-69]; however, little is known about its mechanical impact as imparting rigidness in phospholipid membranes, particularly in the biologically relevant packing status i.e., the hypothetical liquid ordered phase at high (physiological) temperature. In a recent study using NSE with large unilamellar vesicles (LUVs), we reported experimental data on the impact of the *escin*-DMPC molecular interaction in the bending stiffness of highly curved bilayer membranes [70]. From the analysis of the NSE-relaxation times performed in the molecular scale we inferred an *escin-*dependent membrane stiffening as imparted on the fluid phase well above the melting transition of DMPC (*T* ≫ *T*_*m*_ = 23.6 °C). Here we have performed a deeper rheological assessment on the mechanics of the flat membrane configuration in the temperature regimes considered, such as hybrid plates of finite thickness organized behind the Langmuir DMPC monolayers at cohesive interaction with penetrating escin (see Section 3.3; Fig. 3). We measured the rheological response to planar shear as corresponding to in plane reorganizations of the membrane molecules plus the out-of-plane relocation of escin produced under surface deformation (see Experimental; Sections 2.4-2.5).

*Hookean regime: interaction energies*. Figure 5 shows the two rheological scenarios explored either at low temperature (solidlike: blue symbols; upper A panels), or at high physiological-like temperature (fluidlike: red symbols; lower B panels). For the highly rigid escin monolayers the stress-strain plots exhibit nonlinear softening above a yield point detected at very small strains (about *γ* ≈ 1%; see Fig. 5; left panels). The solidlike character of the monophasic escin monolayers is evidenced as an extremely high value of the linear rigidity modulus detected at both temperatures (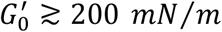; central panels). Such high values of the shear rigidness actually correspond to the strong lateral cohesion energy as above inferred from the surface adsorption energetics (see Section 3.4). In effect, the effective changes in rigidness are expected describing the internal membrane cohesion as the elasticity coupling between the two orthogonal uniaxial deformations i.e., the transverse surface insertion of escin and its longitudinal reorganization [7]. Consequently, the effective shear rigidity must be observed as an apparent surface modulus arising from the membrane cohesion as defined per molecular area i.e.,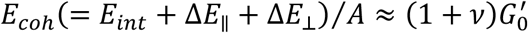(being *v* the Poisson ratio [63]; see Section 3.5). For pure escin monolayers, we have assumed *A*_*esc*_ ≈ 0.9 *nm*^2^ and *E*_*int*_ ≈ 35*k*_*B*_*T* (see Table I), and Δ*E*_∥_ + Δ*E*_⊥_ ≈ 24*k*_*B*_*T* (see Table II), so that we can estimate the surface rigidness as an effective elasticity modulus for the solidlike status i.e., 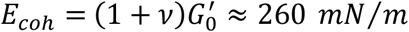 in qualitative agreement with the experimental measurements of shear modulus (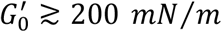 see Fig. 5). Hence, the Poisson ratio is estimated for these bare escin monolayers at *v* ≲ 0.2, as compatible with an extremely high rigidity [10,63]. The solidlike character of these adsorption monolayers of water-soluble escin was further evidenced with respect to the fluidlike behavior of the Langmuir monolayers of the water-insoluble DMPC phospholipid (with apparent values of shear rigidity below the experimental uncertainty; see Fig. 5; central panels).

**Figure 5.**
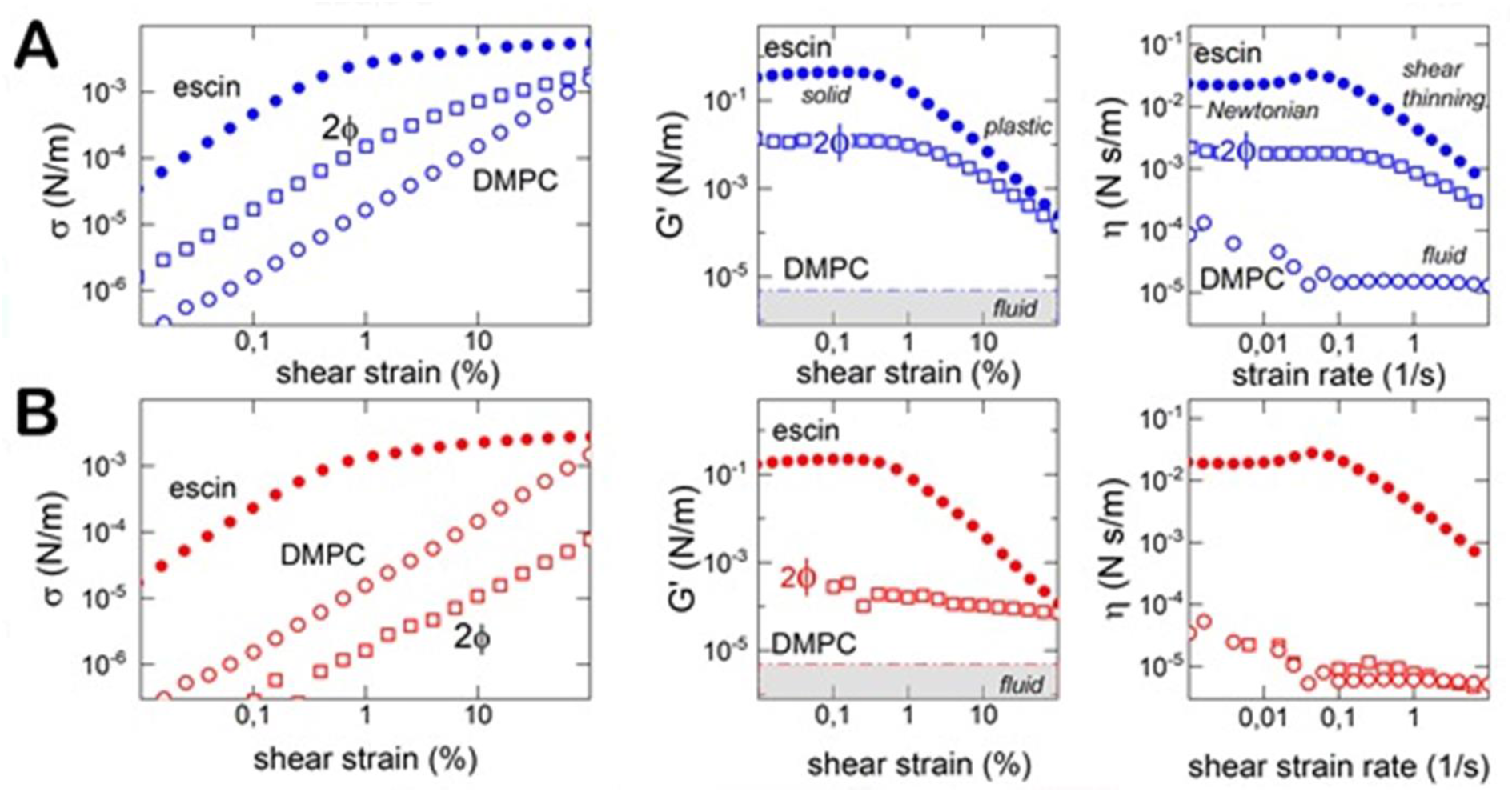
Surface shear rheology of escin / DMPC monolayers. (at the biological reference status *π* = 30 *mN*/*m*). We show comparative graphs of pure escin (monophasic 1ϕ), pure DMPC (1ϕ), and escin interacting with DMPC as penetrating at high insertion degree from the aqueous subphase at concentration 0.1 mM (biphasic 2ϕ); at 4ºC (blue symbols), and 38ºC (red symbols). **A) Stress-strain** plot at 1 Hz. **B) Shear rigidity modulus** as a function of shear strain. **C) Dynamic shear viscosity** as a function of shear strain rate. The bare escin monolayer behaves like a solid, whilst the DMPC monolayer is typically fluid. The mechanical interaction of both components in the escin-penetrated monolayer upon shear depends on the temperature. At low temperature (4ºC) they behave like a soft solid while at high temperature (38ºC), they do like a viscous fluid.

### 3.7 Hydrostatic compression: membrane dilatability and interdomain interaction

*Compression modulus: microscopic dilatability*. Additional to the surface shearing experiments in linear regime, we also performed mechanical experiments probing the hydrostatic compressibility of the DMPC monolayers spread over an escin-containing subphase [10]. Figure 6 shows the experimental measurements performed in the physiologically relevant states of escin / DMPC interaction at high temperature compatible with the liquid disordered molecular status of the phospholipid appeared always monophasic as per entropic dominance. These biomimetic systems of high temperature were compared with the respective enthalpic statuses of high compaction compatible with escin adhesivity appeared, either monophasic for pure escin, or biphasic after the mixing interaction with the liquid ordered phase of DMPC (at low temperature). The compression isotherms plotted in Fig. 6A reveal a dataset with a very similar expanded-like behavior. Indeed, the surface pressure (*π*) monotonically increases from the initial state up to collapse, except for pure escin monolayers and the low temperature statuses of the mixed DMPC/escin monolayers in which escin adsorption is the energetically more favorable at the larger surface areas (i.e., −*E*_*ads*_ = *π*_0_(*A* − *A*_*PC*_); see also Fig. 4B;). In addition, Figure 6B shows the calculated values of the hydrostatic modulus of compressibility *K* = −*A*(*∂π*/*∂A*), which describes the surface response to uniform changes of lateral pressure [10,63]. Interestingly, the nearly conserved bulk modulus of the hybrid DMPC / escin monolayers exhibits a “regulatory” plateau at the bilayer-equivalent packing pressure (*π*_*b*_); this is *K*_*b*_ ≈ 60 *mN*/*m* at *π*_*b*_ ≈ 30 *mN*/*m* (see caption in Fig. 6 for details). At higher compacting pressure (*π* > *π*_*b*_), however, the modulus of lateral compressibility experiences further increases, especially for the very compact systems barely composed by pure escin and mixed with the liquid ordered phase of DMPC. The maximal compression modulus is identified in the pure escin monolayers *K*_*b*_ ≪ *K*_*max*_ ≲ 200 *mN*/*m* at *π*_*max*_ ≳ 40 *mN*/*m* (corresponding to *π*_*ins*_ → 0, the reference state adopted for estimating the energies in Table I and II; see Fig. 3). Indeed, we estimate the cohesion energy of escin under barely adsorption as 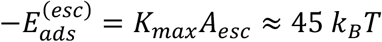 clearly exceeding the longitudinal interaction energy identified for area reorganization i.e., *E*_*coh*_ ≫ Δ*E*_⊥_ (see Table II). Consequently, the effective hydrostatic compressibility plotted in Fig. 5B must be regarded only as a partial component to the global membrane rigidness, namely, a total dilatational surface modulus including compression and shear components; this is *ε* = *K* + *G* ≈ *E*_*coh*_/*A* ≈ (Δ*E*_∥_ + Δ*E*_⊥_ + *E*_*int*_)/*A* [63]. For the considered biomimetic monolayers at high temperature, by taking structural data (*A* ≈ 0.9 *nm*^2^ and Δ*E*_∥_ ≈ 20*k*_*B*_*T*; see Table II), and assuming the Poisson ratio above estimated for the shear measurements (*v* ≈ 0.2), we predict the effective compression modulus at *K*_*b*_ ≈ (1 − 2*v*)*ε* ≈ 60 *mN*/*m*, in qualitative agreement with our experiments (see Fig. 6B). To summarize the impact of escin at mechanical equivalence with a highly rigid material, for solidlike escin-rich membranes interacting with lipids, we have found constitutive properties recapitulated in the surface dilatational rigidity *ε*_*S*_ = *K* + *G* (for 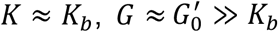 and *v* ≈ 0.2), however, for fluidlike DMPC -rich monolayers we found a similar finite compressibility (*K* ≈ *K*_*b*_), but a negligeable shear stiffness 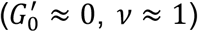 which give rise to a lower dilatational rigidness i.e., *ε*_*F*_ = *K* ≪ *ε*_*S*_. In the opposite case of solidlike monolayers of pure escin, we have found the maximal values of both, the shear rigidity 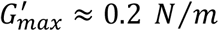 (Fig. 5B), and the compression modulus *K*_*max*_ ≈ 0.2 *N*/*m* (Fig. 6B); hence the dilatational modulus *ε*_*esc*_ ≈ 0.4 *N*/*m*, the Poisson ratio *v*_*esc*_ = (*G* − *K*)/(*G* + *K*) ≈ 0 [63], and the resulting global cohesive energy *E*_*coh*_ = *εA* ≈ 90 *k*_*B*_*T*.

**Figure 6.**
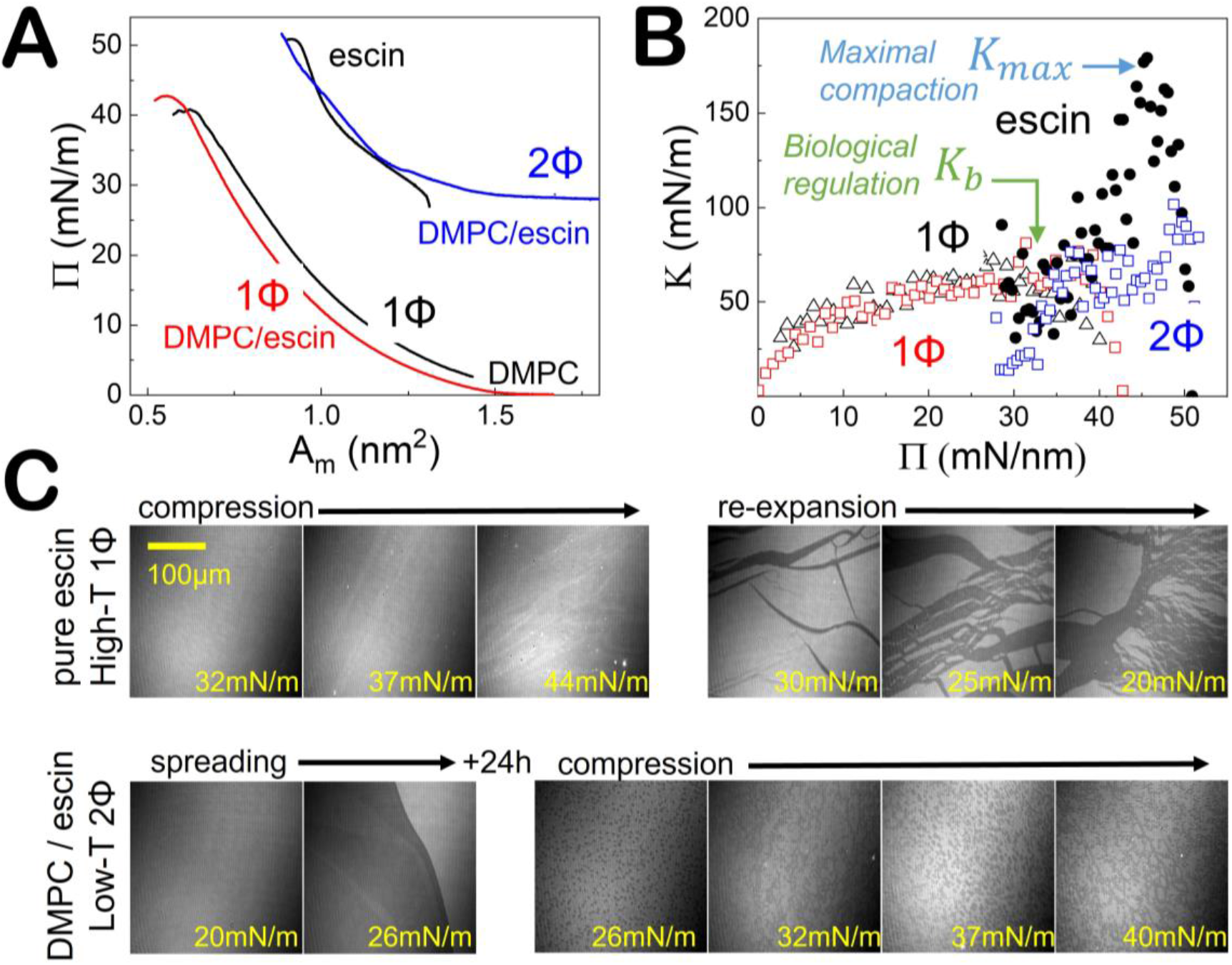
A) Experimental isotherms for uniform lateral compression of escin / DMPC monolayers. (*π* − *A* plots recorded under continuous and uniform change of surface area at the physiological temperature (*T* = 38°*C*); legends included in the image). At low temperature (*T* = 4°*C*), the very rigid monolayer of pure escin resulted extremely brittle under compression being thus impossible an experimental reproducibility in determining the correspondingly unstable compression isotherm. Instead, we have included the low temperature isotherm of the mixed DMPC / escin monolayers as appearing stable in the biphasic state (see Fig. 3). **B) Hydrostatic compression modulus** (i.e., *K* = −*A*(*∂π*/*∂A*)*T* calculated as the numerical derivative of the smoothed isotherms). *Legend: High-temperature states)* monophasic liquid disordered DMPC (1ϕ / L_D_ - black hollow triangles); biphasic escin/DMPC (2ϕ / L_O_/L_D_ coexistence - red hollow squares); monophasic pure escin (1ϕ / solid - black solid circles). *Low-temperature states)* biphasic liquid ordered DMPC / escin (2ϕ / L_O_ -blue hollow squares). Two particular states of biological significance for escin insertion are tagged (corresponding to the high temperature status): *maximal escin compaction* (*Kmax* ≈ 180 − 200 *mN*/*m* at *π* ≈ 40 − 50 *mN*/*m*, corresponding to the virtual state of no insertion pressure *πins* → 0); *bilayer-equivalent state of biological regulation* (*Kb* ≈ 60 *mN*/*m* at *π* ≈ *πb* ≈ 30 *mN*/*m*, corresponding to the virtual bilayer pressure in the liquid ordered structural status of the fluid biological membranes). **C) Monolayer morphologies as determined by BAM under hydrostatic in-plane compression** (surface pressures indicated in yellow): *Upper filmstrip)* solidlike monolayers of pure escin under compression at high temperature before collapse (*πcol* ≈ 52 *mN*/*m*); the further re-expansion causes evident fractures in the solid plates that compose the uniform escin monolayer. *Lower filmstrip)* Phase-separated monolayers of DMPC spread at high temperature on a sub-micellar escin solution (0.1 mM). The monolayers were left overnight to keep reaching the adsorption equilibrium. After 24h, they exhibit the escin-rich crystallite domains already discussed to raft in liquid disordered phase of DMPC (see Fig. 3). The resulting biphasic monolayers were BAM-monitorized under uniform compression from the initial insertion pressure practically till collapse. The mesoscopic bi-phasic superstructure remains progressively crowded as an interacting network of discretely percolated domains composed by the rigid escin (dark domains of aglycone compacted moieties shorter than DMPC), as floating in the continuous phase of the molten phospholipid (appeared brighter as corresponding to longer molecules than escin).

Such parametric constitutive dataset ranks the surfactant β-escin top in the scale of surface cohesivity in fluid interfaces and endows the record of surface rigidness for the uniform (monophasic) adsorption monolayers of a single soluble compound [40].

#### Interdomain interaction: mesoscopic fluidity

Nevertheless, we must keep in mind that the hybrid DMPC / escin system can eventually stay in a heterogeneous biphasic state with solid escin-rich domains rafting the continuous DMPC-rich phase (see Fig. 3). They can eventually lead additional rigidities with a dominant shear rigidity for the highest escin insertions. Hence, an intermediate mechanical impact is observed in these 2Φ-statuses, especially at low temperature (see Fig. 5; central panels). Hence, we visualize the global rigidness as a mesoscopic equivalent *ε*_*eq*_ = *ε*_*eq*_(*ε*_*S*_, *ε*_*F*_, *ε*_*dom*_), including the crowding interaction between the solidlike domains (*ε*_*D*_) [7]. Because a heterogeneous monolayer (raft-like) actually couples the phases’ rigidities in parallel, we predict 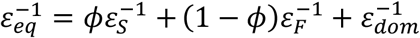(being *ϕ* the domain fraction [64]). As far as the solidlike escin-rich domains interact each other with a remanent 2F-rigidness, in ordered states we thus expect 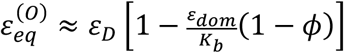for instance at high domain fractions (as experimentally observed at low temperatures *ϕ* → 1; see Fig. 3). This formula shows how much influential are the lateral interactions between domains in the effective rigidness of an ordered mosaic membrane i.e., 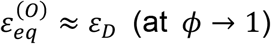 However, for the fluid liquid disordered phase 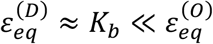 By looking at the experimental data in Figs. 5 and 6, in the highly ordered state at low temperature, we estimate this quantity as the interdomain interaction energy 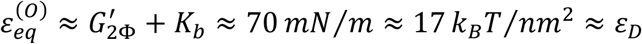 (as calculated per molecule). However, in the highly disordered state at high temperature, we have estimated 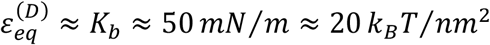 as corresponding to the laterally homogeneous interaction between DMPC and escin at the physiological bilayer state i.e., Δ*E*_∥_ ≈ *K*_*b*_. These mechanical estimations are in qualitative agreement with our previous experimental calculations from the kinetic measurements (see Table I and II), and with more sophisticated theoretical estimates inferred for heterogeneous membranes [7,10,28,33]. Yet being elastically Hookean as corresponding to the relatively low interaction energies put into play at the small deformations considered, all the monolayers studied in this linear, weakly dissipative, regime undergo Newtonian viscous flow at correspondingly low deformation rates (see Fig. 5: right panels).

### 3.8 Nonlinear shear regime: soft solid behavior

The experimental data of shear rheology in Fig. 5 showed an evident nonlinear softening behavior, clearly evidenced for the pure-escin monolayers with a dissipative origin, first as a plastic plateau above a yield stress (left panels), and second, as a decrease of the apparent shear modulus being the consequence of the stress softening above the yield point 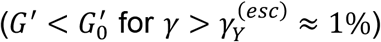.

The results in Fig. 5 also showed the shear viscosity of pure-escin monolayers with values significantly lower than the shear stiffness (*η*_0_ ≥ 0.01 *N s*/*m* in both temperature regimes; Fig. 5; right panels); in the Newtonian regime the dynamic viscosity appears as the linear response to frictional losses at low shear rates below the yield point i.e., 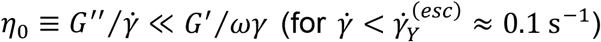 Arguably, the surface friction primarily arises from the dissipation of the cohesion energy that retains the surface molecules adsorbed longer than the diffusion time i.e., *E*_*ads*_ ⇒ Δ*E*_⊥_ ≈ *G*^″^*A* [6,7,12]; by taking the kinetic data measured for pure-escin monolayers in the high compaction low-temperature status i.e., *E*_*ads*_ ⇒ −Δ*E*_⊥_ ≈ 12*k*_*B*_*T*, and *τ*_*D*_ ≈ 10^2^ *s*, by taking *δA*_*esc*_ ≈ 0.3 *nm*^2^ (see Fig. 3 and Table II in Section 3.4; for *A* ⇒ *A*_*esc*_ ≈ 0.9 *nm*^2^ [39,42]), and assuming a diffusional strain rate well within the Newtonian regime 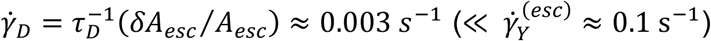we estimate in this regime the highest frictional losses as ω*η*_0_ ≈ *G*^″^ ≥ Δ*E*_⊥_/*A*_*esc*_ ≈ 10 *mN*/*m*, corresponding to the shear viscosity *η*_0_ ≈ *G*^″^/ω ≥ 0.01 *N s*/*m* (in agreement with the experimental data at ω = 1 *Hz*). Above the diffusional yield point 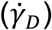 a shear thinning follows characteristic for a soft viscoelastic solid (*η* ≪ *η*_0_ for 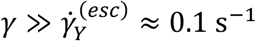 (see Fig. 5; right panels). The stress softened monolayers display apparent values of the shear modulus comparatively higher than found for the loss modulus (i.e., 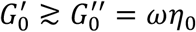 for soft solids), hence defining the viscoelastic status of the monophasic (pure)-escin monolayer to be predominantly a soft solid under weakly-perturbed mechanical equilibrium. This very particular rheological behavior assigns the bare escin monolayers with the mechanical characteristics typical of a two-dimensionally viscoelastic material able to support high shear stress *–a soft solid exhibiting indeed one of the highest values of shear rigidity reported to date for surface films supported on liquid surfaces* [40]. For the bare DMPC monolayers, however, the liquid-crystalline lipid arrangement undergoes linear deformation over the entire range of strains (Fig. 5; left panels), becoming hence able to deform fluidlike (i.e., *G*^′^ = 0; central panels). As expected, the shear viscosities of the DMPC monolayers are meaningfully smaller than found for escin monolayers (left panels). Therefore, the liquid-crystalline mesophases of the bare DMPC monolayers are considered to behave essentially fluidlike in mechanical terms, akin a viscous fluid, not only in the high temperature liquid-disordered (L_D_)-phase, but also in the low temperature liquid-ordered (L_O_)-phase (i.e., *G*^″^ = ω*η* ≫ *G*^′^ = 0 for fluids [7]).

For the binary monolayers of liquidlike DMPC as transversely penetrated by solidlike escin, the biphasic monolayer is observed to produce an intermediate rheological scenario (see Fig. 3). At low temperature we still detect in the linear part of the stress-strain plot a dominance of the solid escin domains (in both temperature regimes, we detected 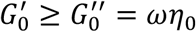 see Fig. 5). As due to the mesoscopic ordering induced by the solid escin-rich domains into the continuous Lo-phase of DMPC at low temperature (as recording nonlinear shear responses at *T* = 4°*C*), we indeed detected 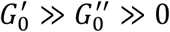 in the widely explored range of deformations. We also detect a plastic yield now appearing shifted up to shear deformations higher than observed for pure escin 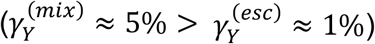 Such a low-temperature DMPC-rich phase seems extrinsically rigidified by integrating escin from the subphase (similarly to a confined liquid-crystal). However, it could be unable to retain shear stress anymore above the yield point (similarly to a high viscosity liquid-crystal under plastic flow). Differently, the high temperature (liquidlike) mesophases of the binary DMPC/escin system do behave rheologically similar to the bare DMPC monolayer (as recording shear responses at 38ºC, we detected 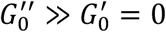 see Fig. 5). As far as the disordered DMPC L_O_-phase can be intrinsically penetrated by escin at high temperatures (above *T*_*m*_ ≈ 23 °*C*), the solid character of the escin-rich domains of the biphasic DMPC-escin systems persists dominant only at low temperature (at *T* ≪ *T*_*m*_), and low deformation strain 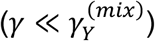 Contrarily, the phase separated DMPC/escin monolayer behaves essentially fluidlike at high temperatures (*T* ≫ *T*_*m*_), and high deformations 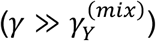 akin a fragile glassy material. In summary, whereas the rigidity modulus is practically null (or undetectable) in DMPC-rich monolayers, as expected for liquidlike phospholipid membranes above the melting transition, the adsorption association with underlying solid escin imparts enough shear rigidity to make the hybrid DMPC / escin membranes behaving likewise a soft solid material.

### 3.9 Surface dynamics in frequency domain: soft glassy rheology

In rheological terms, a typical soft solid is dynamically characterized by a weak dependence of the viscoelastic moduli on the frequency of the deformation; particularly, in the Hookean (linear) regime the rigidity modulus and the slightly lower loss modulus are expected to scale with a similar power law as *G*^′^∼*G*^″^∼ω^*x*^ (with *x* ≪ 1), a behavior widely known as *soft glassy rheology*, or briefly SGR [71,72]. To further evaluate the dynamic behavior of escin-containing membranes (in Langmuir monolayers spread at the A/W-interface), we have analyzed the frequency dependence of the viscoelastic moduli in the structural scenarios above described.

Figure 7 shows the experimental results as comparing the rheological behavior of the binary escin / DMPC monolayers with the single component systems. In the case of pure escin monolayers the viscoelastic moduli are found with similar absolute values in both, high-T and low-T states, and frequency dependencies compatible with SGR behavior (Fig. 7; left panels). Remarkably, we obtained extremal values of shear rigidity (ca. 100 mN/m), even higher than found for the extremely rigid ceramide molecule [9-11], an essential messenger involved in cell apoptosis [13,14]. On the one hand, we detect the solidlike escin monolayers behaving SGR-like with a shear modulus displaying a weak power-law dependency (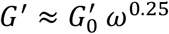 under amplitude 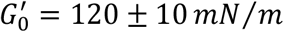), found even weaker for the loss modulus (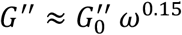 under amplitude 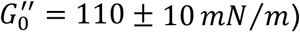). On the other hand, we detect the liquidlike DMPC monolayers behaving in a purely Newtonian fluid scenario at vanishing rigidness both at low and high temperature (i.e., *G*^′^ ≈ 0 and *G*^″^ ≈ ω*η*_0_), corresponding to a time-independent value of the dynamic viscosity (*η*_0_). We indeed measured for DMPC a shear viscosity decreasing with increasing temperature as corresponding to Newtonian fluids (see Fig. 7; right panels; bottom). For the binary DMPC/escin monolayers, however, a completely different behavior was observed (Fig. 7; central panels). When the escin is inserted into the biphasic DMPC monolayer at low temperature (2Φ), a high shear rigidness remains 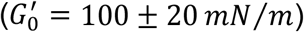 although displaying a stronger frequency dependency 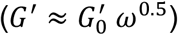 significantly higher than expected for canonical SGR behavior in a rigid solid (*x* ≈ 0). A similar behavior is found at low temperature for the loss modulus (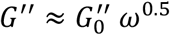 with amplitude 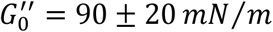). Because 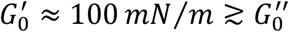 in this solidlike phase - separated state (2Φ, low-T), we still assign it with SGR-like behavior typical for a soft solid. At high temperature for the monophasic liquidlike phase (1Φ, high-T), both, the elastic and loss modulus have finite absolute values albeit relatively inverted as corresponding to a predominantly fluid viscoelastic material 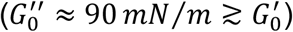 A completely different dynamics is observed breaking out the solidlike SGR behavior typical for shear deformations at low frequencies (ω < ω_*c*_ ≈ 1 *Hz*), up to liquidlike behavior observed at higher frequencies (ω > ω_*c*_; see Fig. 6; central panels). Such dynamic bimodality happens through a characteristic diffusive crossover 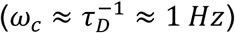 as being compatible with the observed dual diffusion / reorganization kinetics (see Fig. 4). At lower frequencies (ω ≪ ω_*c*_), the slowly deformed DMPC / escin monolayer behaves likewise a soft glassy solid as far as the much faster diffusive exchanges with the bulk are blocked. At higher frequencies (ω ≫ ω_*c*_), however, the monolayer deformed quickly and behaves likewise a regular fluid. It is a viscoelastic material able to relax stresses through of the dynamically congruent adsorption / desorption exchanges with the bulk. Therefore, the rheological dynamic bimodality here revealed for the binary DMPC / escin monolayers could be plausibly connected with the dual phase behavior above discussed in structural terms. Because these molecules are able to partially segregate into two phases of model systems at dependence of temperature, one with a higher phospholipid content behaving liquidlike, and the other with a higher escin content behaving solidlike, such mechanical duality enables the emergence for a complex rheology with a probable biological impact as due to rigidness in mosaic membranes as receiving natural saponins e.g., from foods, or in some therapies.

**Figure 7.**
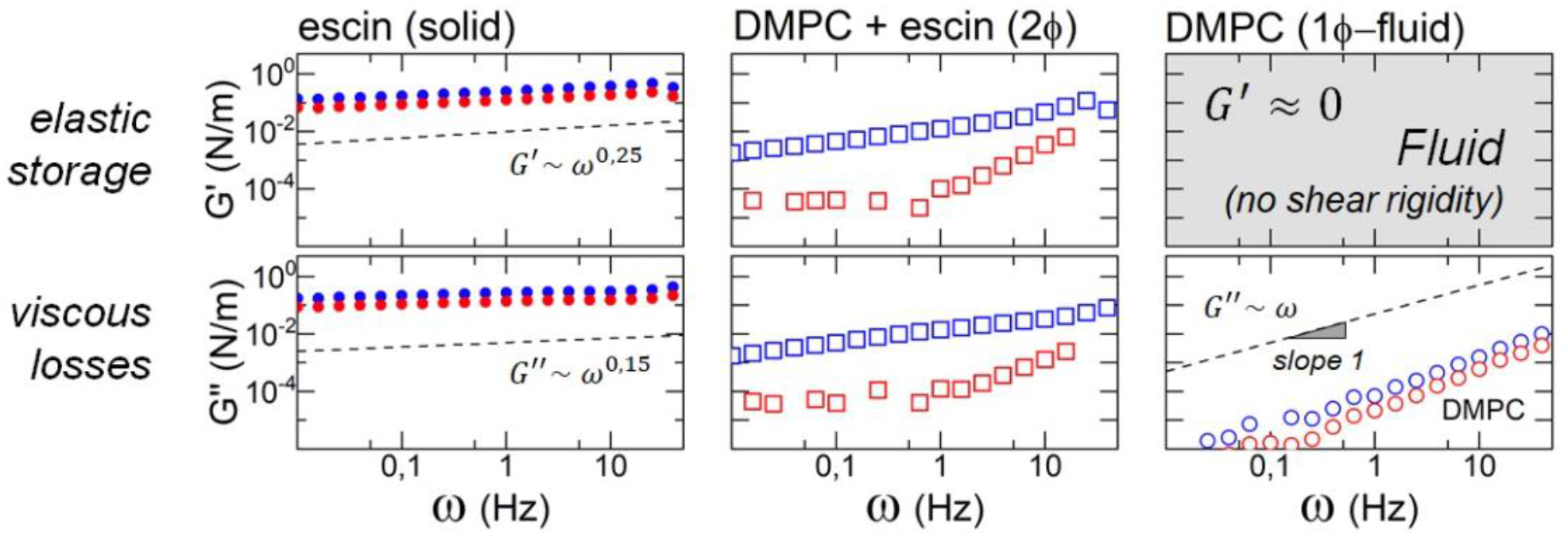
Frequency dependence of the surface shear rheology of escin / DMPC monolayers in the linear regime of shear deformation (elastic storage: upper panels; viscous losses; lower panels). We show comparative graphs of pure escin (monophasic 1ϕ; left panels), escin interacting with DMPC as penetrating from the aqueous subphase at concentration 0.1 mM (biphasic 2ϕ; central panels), and pure DMPC (1ϕ; right panels); at 4ºC (blue symbols), and 38ºC (red symbols).

## 4. Discussion

In this study we have addressed the role of escin in membrane mechanics regulation. The natural saponin acts as an external complexer of membrane mosaicity widely used, for instance, in pharmacological treatments of chronic venous insufficiency (CVI). The amphiphilic character of escin confers this molecule with the dual property to either self-aggregate extrinsically into rigid solid phases of biological membranes with a natural mosaicity, or to intrinsically associate with other membrane components that produce rigidification as interacting with structural elements determining their modified mosaicity. The experiments performed with model systems demonstrate a membrane insertion of the escin complexer into either the longitudinal in-plane integration mode, or the transverse out-of-plane penetration pathways which impact differently the effective rheology of the escin-modified membrane. Weather escin is “congenitally” incorporated to the membrane *ab initio* (longitudinal insertion), the system raises immediately the mechanical equilibrium as a soft glassy solid with a phase and mechanical behavior that depends exclusively on temperature and composition. However, when escin is extrinsically incorporated from the adjacent subphase *ad finem* (transverse insertion), a bimodal adsorption mechanism governed by bulk diffusion / membrane reorganization emerges, which in turn is limited by compositional and thermal conditions. In particular, when adsorbed from the bulk on a phospholipid membrane surface, first a thin layer of saponin surrounds the membrane from its outer side, to later form the integrated escin/lipid membrane. During this diffusional rearrangement a rigidification of the membrane is promoted, limiting its overall elasticity, and increasing the lateral viscosity in dependence on its composition and temperature. Our complementary measurements of compression (*K*), and shear (*G*) rigidness have evidenced the key role of the dilatational modulus (*ε* = *K* + *G*) as a global measurement of molecular compactness as leading the necessary membrane rigidity for functional stability.

Whilst the ordering rigidness has been evidenced at low temperature to govern the escin / phospholipid membranes into solidlike heterogenous arrangements of membrane rafts, at high temperature a mechanical softening concomitant with a more viscoelastic behavior has been observed to dominate in the more homogeneous fluid membranes. The surface shear rigidity and the hydrostatic compressibility have been both reveled controlled by the lateral compaction energy (−Δ*E*_∥_ ≈ 20 − 12*k*_*B*_*T* per molecule), whereas the shear viscosity governing the frictional losses of the membrane increased with the strength of the transverse attractive potential between the supporting surface and underlying escin (−Δ*E*_⊥_ ≈ 12 *k*_*B*_*T* per molecule). In the case of escin-rich membranes, the high longitudinal attraction between the aglycones makes the viscosity very high and dependent on the higher shear rates. Above the diffusional frequency a shear thinning appears in which the viscosity decreases as due to the transversal ejection of escin. For pure DMPC, however, the Newtonian membrane viscosity is independent of the shear rate as being characteristic for the high fluidity of unsaturated phospholipids compared to the high thickening of pure escin. For the mixed DMPC / escin case resembling a real composite biomembrane, at low temperature, the monolayer exhibits a pseudoplastic solidlike behavior compatible with a soft glassy rheology, whilst at high temperature, a Newtonian fluidlike behavior emerges and the system flows likewise the natural phospholipidic membrane. After the stiffening phase owned by the cohesive properties of the membrane assembly, escin begins to nucleate as an aggregated phase of aglycones moieties interacting by hydrophobicity within the lipid bilayer. This nucleation seems to be favored by the pre-existing ordering of the lipids, i.e., the greater the order, the greater the capacity for scaffolded insertion and further reorganization of the saponin in the lipid assembly to promote higher rigidness under regulated frictional losses.

Once the escin molecules are an integral part of the membrane assembly, which in our study would represent a longitudinal insertion similar to natural (congenital) assembly of biological membranes (*E*_*int*_ > Δ*E*_∥_ ≫ Δ*E*_⊥_), they tend to associate with the phospholipids to laterally segregate into two heterogeneous phases –similarly to cholesterol-driven formation of rafts in eukaryote biomembranes [2]. The interaction energies that dominate the raft assembly over the relevant micro- and meo-scales is clearly dominated by the strong surface adhesivity of escin not only in the form of microscopic cohesion between molecular aglycone moieties but also as a cohesivity between domains. This is a multiscale kind of membrane mosaic ordering also favored by the parental lipid order but leading to emergent properties as encoded by the multiscale cohesive energy *E*_*coh*_ = *εA* = (*K* + *G*)*A* [1-5]; for β-escin *E*_*coh*_(= *E*_*int*_ + Δ*E*_∥_ + Δ*E*_⊥_) = *εA* ≈ 90 *k*_*B*_*T*, this enormous cohesive energy has been here estimated being top in the ranking of soluble surfactants known, thus leading the highest surface rigidity able to support hydrodynamic crystallization in water surfaces, for instance [40]. Consequently, when the intrinsic lipid ordering induced by escin becomes higher, the lateral freedom of movement within the escin-rich membrane domains crowded, and the transverse diffusivity of escin is also strongly hindered, the escin-containing membrane becomes then the most rigid. However, the natural segregation into two mosaic fluid phases should make the rigid membrane to mesoscopically fluidify weather the continuous phase appears more fluid (as composed of molten lipids in the L_D_-state) than the discrete, much more rigid, domains (enriched in escin and ordered lipids). Mesoscopic fluidity emerges then as a tradeoff between the cohesive escin / lipid interactions (enthalpic) playing against the disordering domain interactions (entropic). Such composite mosaic structure imparts greater effective fluidity through of the continuous incompressible phase of liquid-crystalline lipids as occurred in biological membranes. This rheological diaphony leading to mesoscopic fluidity within a heterogeneously rigid membrane system, rather meaning a synergistically dual functional mechanics than a negative antagonism, is not rare to appear in more complex lipid systems endowing the compositional heterogeneity that characterizes the natural fluid mosaicity of real biological membranes. The mechanical diaphony specifically found in the rheology of the saponin / phospholipid interaction could play an important role in the therapeutic efficacy of pharmacological procedures involving the interaction of aglycone moieties with the membrane phospholipids. During the initial stages of treatment, when the saponin associates at the cell surface, the β-escin could promote dual cell rigidness and / or softening at interference with cholesterol and / or other structural mediators of membrane rigidness and fluidity [45,50-52], which could be subsequently corrected when this active molecule becomes inserted into the membrane core. The present results represent a physicochemical insight that opens new avenues of thinking for membrane targeted treatments with saponins based on their mechanical and structural dual mechanics behaving in biological membranes both as solids and liquids.

## Acknowledgments

LHM is contracted by Mar*í*a Zambrano Program from Ministerio de Universidades de España for the attraction of international talent under Next Generation European Union funding (grant CT19/22). The work was supported by the Spanish Ministry of Science and Innovation (MICINN–Agencia Española de Investigación AEI) under grants PID 2019-108391RB-100 and TED2021-132296B-C52 (to FM), and Comunidad de Madrid under grants S2018/NMT-4389 and Y2018/BIO-5207 (to FM). We also acknowledge the financial support of the German Research Foundation DFG Grant HE 2995/7-1 (to TH), and the Open Access Publication Fund of Bielefeld University for the publication costs. This study was also funded by the REACT-EU program PR38-21-28 ANTICIPA-CM, a grant by Comunidad de Madrid and European Union under FEDER program, from European Union in response to COVID-19 pandemics. The funders had no role in the study design, data collection, analysis, preparation of the manuscript, or the decision to publish. We acknowledge Prof. Juan J. Giner-Casares for making available Langmuir troughs and the BAM facility in his laboratory at Universidad de Cordoba, and also for fruitful discussions on the monolayer domain morphologies. LHM gratefully thanks him for hospitality.

## Authors contribution statement

LHM, DHA, MTMR, NC, CD and RG conducted research, provided experimental data, contributed in analyzing data. JAS, TH and FM supervised research and drafted the manuscript. TH and FM supported the search for funding, planning the research, supervised the research, contributed in analyzing data, and wrote the manuscript.

